# Interhelical E@*g*-N@*a* Interactions Modulate Coiled Coil Stability within a *De Novo* Set of Orthogonal Peptide Heterodimers

**DOI:** 10.1101/2023.05.23.541579

**Authors:** Anthony R. Perez, Yumie Lee, Michael E. Colvin, Andrea D. Merg

## Abstract

The designability of orthogonal coiled coil (CC) dimers, which draw on well-established design rules, plays a pivotal role in fueling the development of CCs as synthetically versatile assembly-directing motifs for the fabrication of bionanomaterials. Here, we aim to expand the synthetic CC toolkit through establishing a “minimalistic” set of orthogonal, *de novo* CC peptides that comprise 3.5 heptads in length and a single buried Asn to prescribe dimer formation. The designed sequences display excellent partner fidelity, confirmed via circular dichroism (CD) spectroscopy, and are corroborated *in silico* using molecular dynamics (MD) simulation. Detailed analysis of the MD conformational data highlights the importance of interhelical E@*g*-N@*a* interactions in coordinating an extensive 6-residue hydrogen bonding network that “locks” the interchain Asn-Asn’ contact in place. The enhanced stability imparted to the Asn-Asn’ bond elicits an increase in thermal stability of CCs up to ∼15°C and accounts for significant differences in stability within the collection of similarly designed orthogonal CC pairs. The presented work underlines the utility of MD simulation as a tool for constructing *de novo*, orthogonal CCs, and presents an alternative handle for modulating the stability of orthogonal CCs via tuning the number of interhelical E@*g*-N@*a* contacts. Expansion of CC design rules is a key ingredient for guiding the design and assembly of more complex, intricate CC-based architectures for tackling a variety of challenges within the fields of nanomedicine and bionanotechnology.

## 1. Introduction

As the selected machinery in biology, proteins serve an expansive array of functional roles. Among their many roles, natural proteins mediate a host of chemical transformations, regulate intercellular communications, neutralize pathogens, and serve as integral structural components in connective tissue and the extracellular matrix. The broad utility of proteins undergirds several research fields including protein design, bioengineering, and protein self-assembly, which all aim to broaden and enhance the application scope of proteins for tackling a variety of technological challenges. While naturally occurring proteins represent a logical starting point for the development of protein-based technology, peptides, including polypeptides, have emerged as viable alternatives for replicating the structure and function of proteins.^1, 2^ Peptides exhibit the same chemical diversity as proteins and retain many of their structural features including various secondary structures and folding motifs (*e.g.*, alpha helices, collagen triple helices, coiled coils, etc.). The short length of peptides facilitates the ability to establish sequence-to-structure relationships, which is challenging to elucidate for more complex protein systems. Remarkably, these short amino acid chains still fulfill many of the roles inherent to proteins, including their biological function and assembly (*e.g.*, short RGD peptides facilitate integrin binding).^2, 3^ Moreover, as synthetically tractable biopolymers, peptides, from a synthetic standpoint, are extremely versatile in that an array of non-proteinogenic motifs can be programmed into the sequence design to create peptide-hybrid molecules via inclusion of non-canonical/peptidomimetic residues (*e.g.*, β-amino acids, peptoids, aza-glycine, etc.),^4–8^ oligonucleotides,^9–11^ and small molecules.^12^ As the repertoire of peptide-based chemical transformations expand, the chemical, structural, and functional scope of peptides will continue to broaden, positioning these biomolecules as key components in the development of novel biomaterials with properties and function that extend beyond nature’s capability.

Coiled coil (CC) peptides and polypeptides have emerged as programmable, biosynthetic tectons for the rational design and construction of artificial protein-mimetic systems and architectures.^13–25^ CCs, which consists of two or more alpha helices that intertwine to form a left-handed superhelical coil, are characterized by a seven-residue heptad repeat that is alphabetically assigned *abcdefg* (**Figure 1**). The driving force for association stems from so called “knobs-into-holes” interactions of hydrophobic side chains of residues that comprise the *a* and *d* positions, which constitute the interhelical interface (**Figure 1**).^26^ Complementary charged residues, which typically occupy the *e* and *g* positions, provide additional stability via interhelical salt bridges and assist in defining the helical orientation (parallel vs. antiparallel).^27^

**Figure 1.**
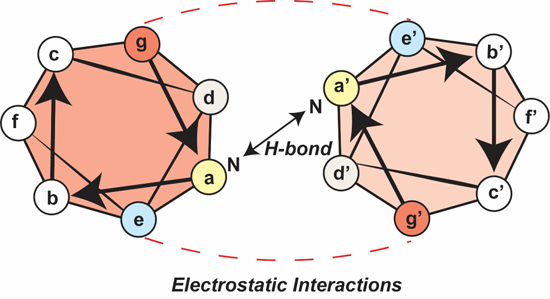
Helical wheel diagram for parallel CC heterodimers showing the N@*a*-N@*a*’ hydrogen bonding interaction and interhelical salt bridges between *g* and *e* positions.

A key feature of CCs is the ability to direct predetermined pairings between CC peptides through rational engineering of the peptide sequence. Decades of research on CCs has yielded a rich set of design rules and considerations,^28–42^ which have allowed for the construction of several collections of orthogonal CC sets.^17, 43–52^ The well-established sequence-to-structure relationship of CCs distinguishes CC-based assemblies from other peptide self-assembly platforms that rely on relatively non-specific and non-programmable interactions. While current working design rules have brought forth inspirational nanoarchitectures, including *de novo* protein polyhedral cages,^13^ the expansion of CC design rules remains a critical step towards increasing the complexity and function of these architectures for potential applications in nanomedicine and nanobiotechnology.^13, 22, 46, 53–55^

Here, we establish a “minimalistic” set of short, orthogonal CC peptides and employ molecular dynamics (MD) simulations to help provide structural explanations for the experimental binding data. We aimed for a CC set that: 1) consists of the minimum length required for forming stable CCs with melting temperature (T_m_) values between 30°C and 60°C (an intermediate range that is suitable for dynamic interactions); 2) displays high partner fidelity; and 3) can be corroborated using MD simulation to facilitate the introduction of new derivative sequences into the CC set. Based on these criteria, we settled on a particular *de novo* sequence design that was first reported by Woolfson *et. al.*^32^ The parent sequence, which comprises ∼3.5 heptads (24 residues), exhibits intermediate stability (T_m_ = 45°C; K_d_ = 5.15 ± 2.05 nM) and, importantly, forms dimers, exclusively. Both features are attributed to the inclusion of a buried polar interaction through placement of Asn in the *a* position. To increase specificity between our designed on-target pairings, Asn residues are incorporated at homologous sites only between sequences of designed pairs. This strategy was similarly employed by Gradišar and Jerala, in which they demonstrated orthogonality of 4-heptad CC peptides comprising two Asn residues.^44^ In our set, however, we decided to introduce a single Asn into the sequence design to counteract the increased instability associated with our shorter CC sequences. Lastly, we highlight the utility of employing MD simulation as a predictor for partner fidelity and as an important tool for disclosing intricate interactions that can be used to refine the CC toolkit and inform the design of future CC-mediated architectures.

## 2. Materials and Methods

All chemical reagents were purchased from Sigma-Aldrich Chemical Co. (St. Louis, MO) or Thermo Fisher Scientific Inc. (Waltham, MA) unless otherwise stated. Fmoc amino acids and Rink amide resin were purchased from CEM (Matthews, NC). UV-vis spectra were collected using a Thermo Scientific Nanodrop One UV-vis spectrophotometer. Matrix-assisted laser desorption ionization time-of-flight (MALDI-TOF) mass spectrometry data were collected using a Bruker Microflex LRF mass spectrometer (positive reflector mode) and using α-cyano-4-hydroxycinnamic (CHCA) as the ionization matrix (1:1 sample to matrix ratio).

### Peptide synthesis and purification

Peptides were prepared using microwave-assisted solid-phase peptide synthesis (SPPS) on a CEM Liberty Blue solid-phase peptide synthesizer and ProTide Rink amide resin (CEM). Fmoc-deprotection was achieved with 4-methylpiperidine (20% v/v) in dimethylformamide (DMF). Coupling of amino acids was achieved via N,N’-Diisopropylcarbodiimide/Oxyma Pure-mediated activation protocols. Peptides were N-terminally acetylated and C-terminally amidated. After synthesis, the peptidyl resins were filtered, rinsed with acetone, and air-dried. The crude peptides were cleaved from the resin for 4 hours at room temperature with a 92.5% trifluoroacetic acid (TFA), 2.5% H_2_O, 2.5% 3,6-dioxa1,8-octane-dithiol, 2.5% triisopropylsilane cleavage solution, precipitated with cold diethyl ether, and centrifuged at 4000 rpm for 10 min at 4 °C. After centrifugation, the supernatants were discarded, and the pellets were dried under vacuum overnight. Crude peptides were purified by high-performance liquid chromatography (HPLC) using an Agilent 1260 Infinity II HPLC instrument equipped with a preparative scale Phenomenex Kinetex XB-C_18_ column (250 x 30 mm, 5 μm, 100 Å). Peptides were eluted with a linear gradient of acetonitrile-water with 0.1% TFA. The target fractions were collected, rotovaped, and lyophilized. Lyophilized peptides were reconstituted in water and quantified by A280 using 1280 M^-1^ cm^-1^ as the molar extinction coefficient of tyrosine. Following quantification, each peptide was aliquoted, lyophilized, and stored at −20 °C.

### Circular dichroism

Circular dichroism (CD) measurements were collected on a Jasco J-1500 spectropolarimeter using a 0.1 mm path length cuvette (Starna Cells, Inc., Atascadero, CA). Three spectra were recorded and averaged from 260 to 190 nm at a scanning rate of 100 nm/min and a bandwidth of 2 nm. CD thermal denaturation experiments were performed by heating from 5 to 80°C (or 90°C) at a rate of 40°C/h. The intensity of the CD signal at 220 nm was recorded as a function of temperature. Lyophilized peptides were reconstituted in 1x PBS (pH 7.4), heated to 90°C for 10 min. and cooled (0.5°C/min) to 5°C. Samples were stored at 5°C prior to CD analysis.

### Thermodynamic parameters

Thermodynamic parameters were obtained using the method described by Marky and Breslauer, and Woolfson *et al*.^32, 56^ Thermodynamic parameters were determined by obtaining full CD spectra at 5 °C intervals from 5 °C to 80 °C (16 data points). The sample holder was heated at a 1°C/min heating rate and the samples were equilibrated for 2 minutes before full spectra was recorded for 200 µM, 100 µM, 50 µM, 20 µM, and 2 µM total peptide concentration samples. The spectra were baseline corrected, and the intensity at 222 nm was extracted from the spectra and plotted as a function of temperature. The points were fit to a Boltzmann curve using Origin. These experiments were done in triplicate, where the melting temperatures were converted to Kelvin, inverted, and averaged. The averages were plotted against the natural logarithm of the total peptide concentration in molarity with the standard deviation used as the y-error bar. Linear-regression was done in Origin to obtain the slope and the y-intercept which were used to calculate `Δ`H and ΔS. For non-self-complementary sequences with a molecularity of 2 (K_eq_ = 4/*C_T_*), the best-fit line has the form:

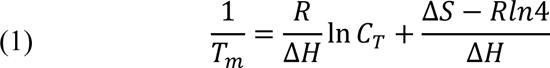

The uncertainties of the slope and y-intercept were used calculate the uncertainty in K_d_ by applying the error propagation formula (2).

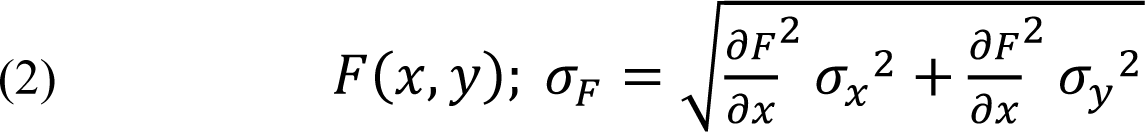

### Size-exclusion Chromatography

Size-exclusion chromatography (SEC) experiments were conducted at 5 °C with a 500 µL sample loop on an ÄKTA fast protein liquid chromatography (FPLC) system, equipped with a Superdex 75 GL 10/300 column. Approximately 500 µL of sample was injected and eluted with 1x PBS (pH 7.4) at a set rate of 0.5 mL/min, and the progress of the separation was monitored at A280 nm. Samples were prepared according to the same protocol used for conducting CD experiments.

### Molecular dynamic simulations

MD simulations were performed using GROMACS version 2022.3.^57^ After short equilibration runs, three replicate 100 ns of MD simulations were run for each structure using the NPT ensemble, the Verlet cutoff scheme and a 2 fs timestep. All bonds to hydrogen were constrained to their equilibrium length using the LINCS algorithm.^58^ Temperature was maintained at 300K using the Bussi et al. thermostat^59^ and pressure at 1 bar using the Parrinello-Rahman barostat.^60^ The simulations were performed using the AMBER99SB-ILDN force field for the protein^61^ and the TIP3P water model.^62^ The total system charge was neutralized by Na^+^ and Cl^-^ ions, and additional ions were added to yield a total ion concentration of 70 mM. The root-mean-square structure deviations (RMSD), root-mean-square fluctuations (RMSF), number of hydrogen bonds, and number of salt bridges were determined using the GROMACS built-in analysis tools. Average values between 25 and 100 ns were obtained using Origin and the statistics gadget on the plotted figure. For each trajectory, snapshots were taken every nanosecond and a custom script using the Selenium package (https://www.selenium.dev/) was used to upload each snapshot to the “Protein interfaces, surfaces and assemblies” service (PDBePISA) at the European Bioinformatics Institute (https://www.ebi.ac.uk/pdbe/pisa/).^63^

## 3. Results and Discussion

Three *de novo* CC pairings (**AA’**, **BB’**, and **CC’**) were designed from six peptide sequences (**A**, **A’**, **B**, **B’**, **C**, and **C’**; **Table 1**). The sequence architecture follows previous *de novo* design rules for forming CC dimers in which β-branched, hydrophobic residues (Ile and Leu), populate the *a* and *d* positions, respectively, and complementary charged residues (Glu and Lys) are placed at the *e* and *g* positions to reinforce partner fidelity via introducing attractive and repulsive electrostatic interactions.^38^ Alanine, a strong alpha helix promoter,^64, 65^ comprises the *b* and *c* positions, and various residues (Glu, Lys, Gln, and Tyr) are incorporated at remaining *f* sites and assist in solubility, ease of purification via HPLC, and quantification via UV-vis spectroscopy. An exception is the placement of Asn at a single *a* site. Although this diminishes the stability of CCs, due to disruption of the interhelical hydrophobic interface, the loss in stability is compensated by the increase in specificity for parallel CC dimers via interchain hydrogen bonding.^30, 41, 44^ The Asn residue is strategically placed in different heptads between designed CC pairs to increase the energy gap between on-target and off-target heterodimers.^44^ Based on these sequences, the thermodynamically favorable CC fold will: 1) have the greatest number of interhelical salt bridges; 2) maximize hydrophobic contact at the interface; and 3) align Asn residues across the helix interface to accommodate enthalpically favorable hydrogen bonds. The partner sequences to **A**, **B**, and **C**, (**A’**, **B’**, and **C’**, respectively) are designed to maximize these enthalpic contributions and contribute to the greatest decrease in the folding free energy. The peptides were synthesized using standard microwave-assisted solid-phase peptide synthesis (SPPS) and purified via HPLC (**Figure S1**). The compositions of the peptides were confirmed via MALDI-TOF mass spectrometry (**Figure S2**).

**Table 1.**
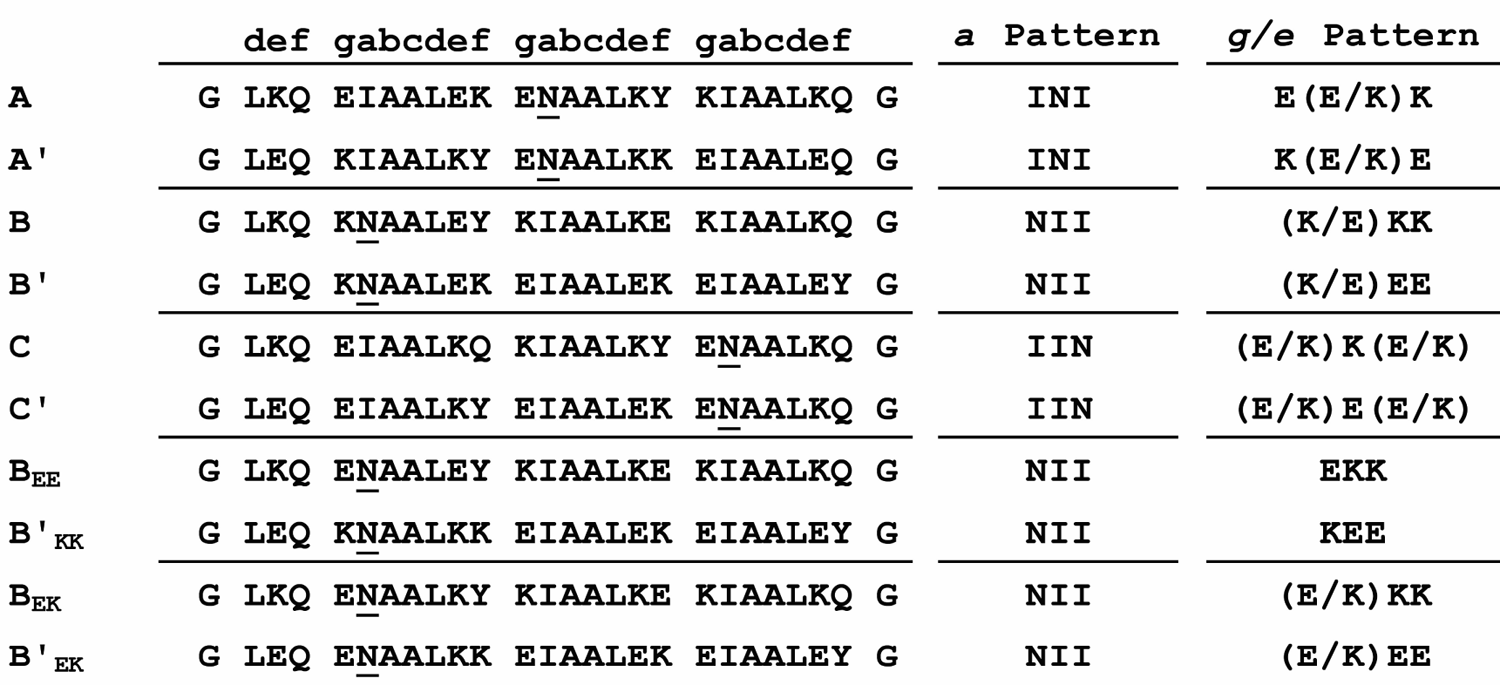
Sequences of *De novo* Peptides that Form Orthogonal Parallel CC Heterodimers

Circular dichroism (CD) spectroscopy was employed to characterize the folding of each individual peptide (**A**, **B**, **C**, **A’**, **B’**, and **C’**) and the three on-target pairings (**AA’**, **BB’**, and **CC’**; **Figure 2**). Prior to CD analysis, all peptides (100 μM) were dissolved in phosphate buffered saline (PBS; pH 7.4), heated to 90°C and cooled to 5°C. CD spectra of the individual peptides display varying degrees of alpha helicity as evidenced by differences in the intensity of the CD signal at 208 and 222 nm (**Figure 2A-C, Table S1**). In all cases, the CD spectra of solutions containing a 1:1 equimolar mixture of the designed on-target peptide pairings exhibit a significant increase in the alpha helical CD signature, and, importantly, differ from CD plots corresponding to the sum of the spectra of the individual peptides (**Figure 2A-C**). Moreover, an increase in the ratio of θ_222_/θ_208_ to values above 1.0 (a range around 1.0 or greater serves as a proxy for the predominance of CC species)^66–68^ is observed when both on-target partners are present in solution, with the exception of the C-series in which **C** and **C’** are relatively structured (**Figure 2A-C**, **Table S1**). To assess the fidelity of the on-target dimers, CD spectra for all off-target pairings were collected (**Figure S3, Table S1**). CD spectra of the mixed peptide solutions closely match with the sum of the individual peptides, which implies a minimal degree of interaction between off-target peptide pairs.

**Figure 2.**
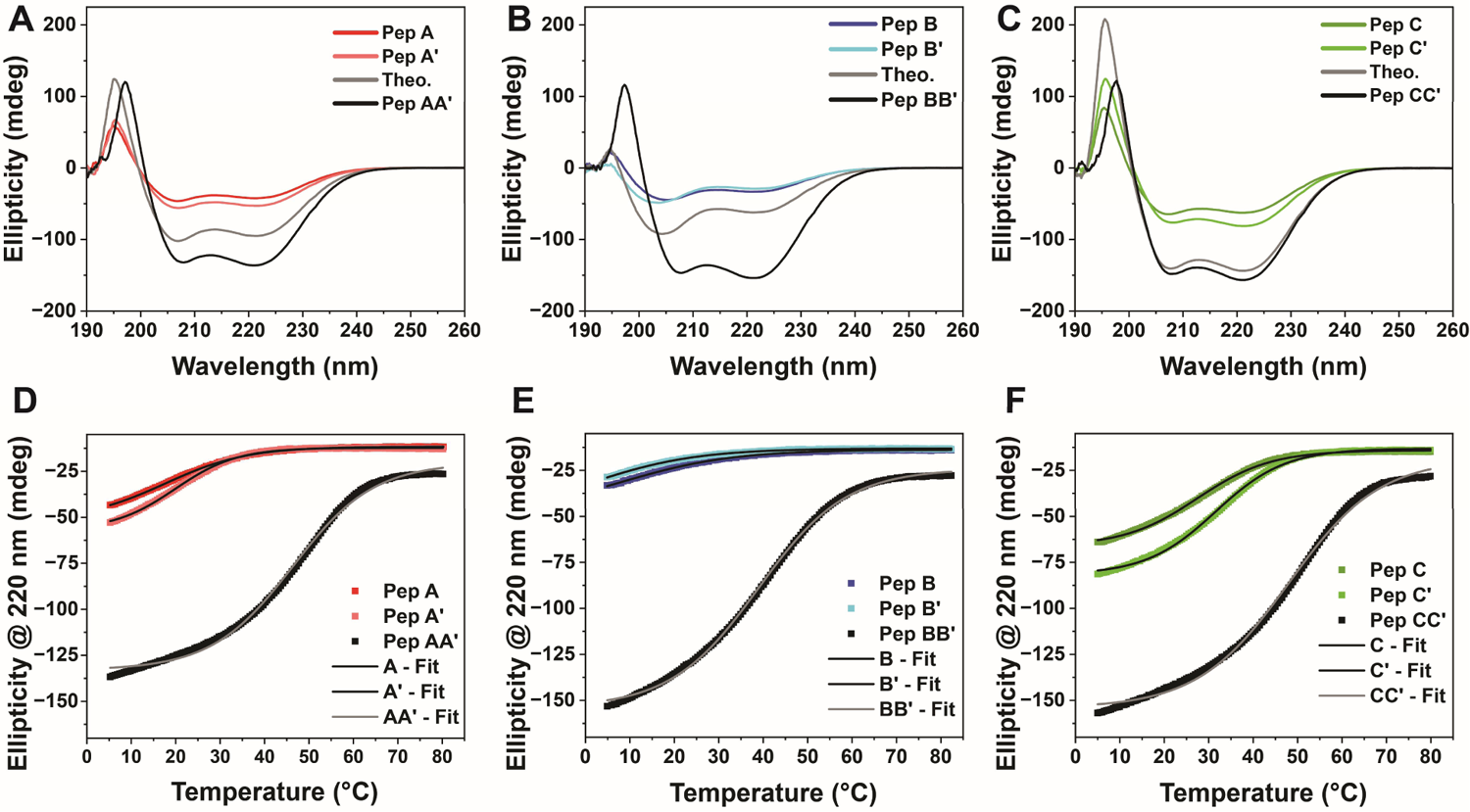
CD spectra and thermal denaturation plots for (**A**) **A**, **A’**, and **AA’**, (**B**) **B**, **B’**, and **BB’**, and (**C**) **C**, **C’**, and **CC’**. Note: the line labeled “Theo.” represents the sum of the plots of the individual peptides. CD thermal melting curves for (**D**) **A**, **A’**, and **AA’**, (**E**) **B**, **B’**, and **BB’**, and (**F**) **C**, **C’**, and **CC’**.

CD thermal denaturation studies were conducted to probe the thermal stability of the CC set, including between off-target pairs (**Figure 2D-F**). T_m_ values for the individual peptides, which were only marginally helical, as assessed from CD, vary from < 5°C (**B’**) to 32°C (**C’**). The wide range of T_m_ values for sequences with nearly identical amino acid compositions underlines the notable role that sequence architecture (*e.g.*, distribution of charges) has on the folding propensity. In contrast, T_m_ values are comparatively higher when complementary designed peptides are mixed in solution (T_m_ = 46°C, 39°C, and 48°C for **AA’**, **BB’**, and **CC’**, respectively; **Figure 2D-F**). The designed heterodimers all exhibit higher thermal stabilities than the off-target pairings (**Figure S4, Table S1**). Altogether, the CD data confirm the orthogonality of the *de novo* CC set. We note, however, from size-exclusion chromatography (SEC), that solutions containing the on-and off-target peptides elute at similar volumes, which suggests the presence of non-specific interactions between off-target sequences (**Figure S5**). Such interactions may explain the relatively high T_m_ and θ_222_/θ_208_ value of **C** and **C’**, and their off-target variants.

Thermodynamic parameters, obtained from measuring T_m_ dependence as a function of peptide concentration, confirm the order of thermodynamic stability (**BB’** < **AA’** < **CC’**), as evidenced by the calculated ΔG values and dissociation constants (K_d_; **Table 2**, **Figure S6, Table S2**). K_d_ values of the on-target designs are higher than that of the parent sequence (K_d_ ≈5 nM). We surmise that the weaker binding interaction of our CC heterodimers stems from the mixed distribution of electrostatic charges along the peptide sequences. The parent sequences are comprised solely of Glu or Lys at the *g* and *e* positions, which contrasts the varied electrostatic pattern that we utilized as a means to introduce positive and negative design elements for creating orthogonal pairings. Prior studies have shown that greater intramolecular electrostatic attraction reduces interchain attraction, thereby lowering the thermal stability of CCs.^50^ The acquired thermodynamic data foreshadow similarities between **AA’** and **CC’**. Both heterodimers have similar changes in enthalpic and entropic contributions to the free energy of binding, which results in similar K_d_ values. On the other hand, **BB’** possesses the highest relative free energy primarily due to having the highest entropic penalty of the three on-target designs. This conforms with CD data, which shows that the **B** and **B’** are least helical, and therefore folding between these monomers must pay a high entropic cost. We next turned to MD simulations to assess the quality of agreement of MD with the experimental data and potentially elucidate the apparent differences between the three on-target dimers.

**Table 2.** Calculated T_m_, ΔH, ΔS, ΔG, and K_d_ Values from CD Thermal Unfolding Curves

| | $T_m$ ($^{\circ}\text{C}$) | $\Delta H$ (kJ/mol) | $\Delta S$ (J/K·mol) | $\Delta G$ (kJ/mol) | Rel $\Delta G$ | $K_d$ (nM) |
| --- | --- | --- | --- | --- | --- | --- |
| AA' | 46.3 | -152.70 | -394.65 | -37.01 | 1.20 | 254.65 ± 0.41 |
| BB' | 38.9 | -176.28 | -485.59 | -33.93 | 4.27 | 899.03 ± 1.48 |
| CC' | 47.9 | -164.72 | -431.58 | -38.21 | 0.00 | 156.42 ± 0.06 |
| B<sub>EE</sub>B'<sub>KK</sub> | 45.4 | -170.40 | -453.68 | -37.41 | 0.80 | 215.86 ± 0.24 |
| B<sub>EK</sub>B'<sub>EK</sub> | 53.0 | -148.07 | -370.50 | -39.46 | -1.25 | 093.02 ± 0.12 |

### MD Simulations Resolve Differences Between On- and Off-Target Dimers

MD simulations were performed on all on-target and off-target heterodimers (15 total pairs). For each pair, 100-ns simulations were carried out at 300 K in triplicate using the GROMACS software package^57^ and the AMBER99SB-ILDN force field.^61^ Initial structures were constructed from point-mutations in Pymol using the PDB of CC-Di obtained from CCBuilder2.0.^31, 69^ Root mean square deviation (RMSD), which measures the structural deviation of backbone atoms from starting positions, for on-target pairs (**AA’**, **BB’**, and **CC’**) display low RMSD values that remain constant throughout the duration of the simulation (**Figure 3A** and **S7**). In contrast, off-target complexes, with the exception of **A’B**, were far more dynamic with greater structural deviations and run-to-run variability, implying that these CC dimers are weakly bound (**Figure S8**). Similarly, the root mean square fluctuation (RMSF), which measures the individual mobility of the C_α_ along the peptide sequence, at the *a* and *d* positions (*i.e.*, residues that constitute the interhelical interface) are comparatively lower for on-target complexes than their off-target counterparts (**Figure 3B** and **S9-10)**. Altogether, RMSD and RMSF analysis indicate that the designed CC pairs maintain tight interhelical contacts that persist throughout the CC interface, unlike the off-target dimers which exhibit greater structural fluctuations and mobility (**Table S3**). These results are consistent with the CD thermal denaturation experiments, in which on-target heterodimers display higher T_m_ values (greater thermal stability) in comparison to the off-target complexes (**Table S1**).

**Figure 3.**
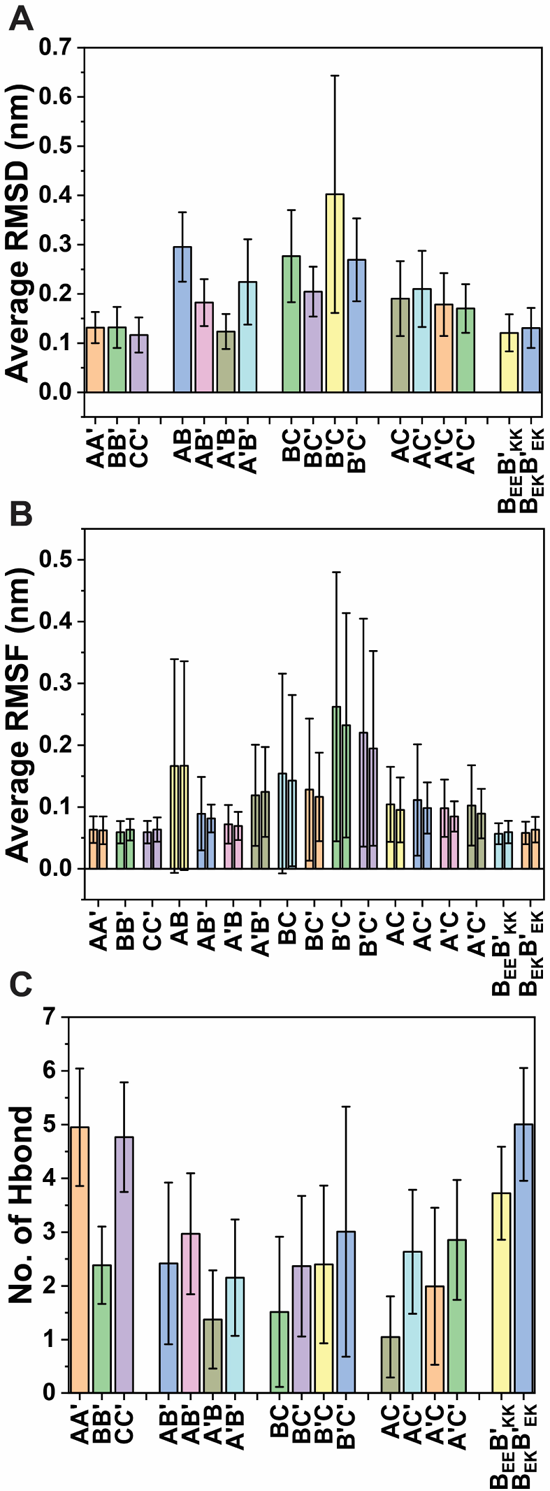
Average (**A**) RMSD, (**B**) RMSF (for residues at the *a* and *d* positions), and (**C**) number of interhelical hydrogen bonds for all 15 simulated pairs. Tabulated values can be found in **Table S3**.

While RMSD and RMSF data correctly distinguish (apart from **A’B**) differences in stability *between* the on-target and off-target variants, both sets of data are unable to adequately recapitulate the finer differences in thermal stability *within* the on-target and off-target variants. In some cases, RMSD and RMSF data accurately resolve relative differences in CC stability between off-target designs (*e.g.*, **A’B** and **BC’** both have the lowest RMSD/RMSF values and highest T_m_ within their group). However, in many cases the RMSD and RMSF data on their own fail to resolve the experimental results, including the notable difference in thermal stability between **BB’** and the other on-target dimers (**AA’** and **CC’**), and especially when comparing off-target dimers between groups (*e.g.*, T_m_ of **BC’** >> T_m_ of **A’B**). We attribute the discrepancy between MD simulation and CD analysis to the large variability that is observed between triplicate runs, especially for off-target pairings, in which standard deviations overlap across the different samples (**Figure 3A,B**). The results highlight the suitability of our relatively short timescale MD simulations to predict stability between on-target and off-target dimers, but also demonstrate its limitation in resolving finer differences between CC dimers that contain few attractive/stable contacts. Alternative MD protocols, at much larger aggregate computational costs, could potentially be employed to improve correlation between experimental and computational results, such as carrying out simulated annealing and/or employing replica exchange molecular dynamics (REMD) simulation to more completely sample configuration space across multiple temperatures, or much longer time scale simulations leading to chain separation or denaturation events.

### Extensive Hydrogen Bonding Network Stabilizes Interhelical Asn-Asn’ Contact

Notable differences emerge between on-target heterodimers when comparing the average number of hydrogen bonds obtained from the MD simulations (**Figure 3C**). While **AA’** and **CC’** average ∼5 interchain hydrogen bonds for the duration of the simulations, which far exceeds the off-target heterodimers, **BB’** possesses on average only ∼2.5 hydrogen bonds (on par with the off-target pairs; **Figure 3C, S11-12**). Unlike RMSD and RMSF, in which **BB’** exhibits similar values to **AA’** and **CC’** (**Figure 3A,B**), the decreased number of interhelical hydrogen bonds observed for **BB’** provided the first possible insight for explaining the comparatively lower T_m_ and higher K_d_ value of **BB’**. This distinct result for **BB’** was further confirmed by analysis of the number of interchain salt bridges, in which **BB’** exhibits fewer salt bridges, on average (0.63 for **BB’** versus 1.03 and 0.89 for **AA’** and **CC’**, respectively; **Figure S13-15**).

In-depth investigation via PDBePISA analysis, which provides detailed information regarding interchain surface contacts including hydrogen bonds and salt bridges, reveals key differences that distinguish **BB’** from the other designed heterodimers. Although the Asn-Asn’ hydrogen bond represents the interhelical contact with the highest percentage of occurrence for all three on-target pairs, the percentage of time during the simulations that this hydrogen bond was present is, on average, lower for **BB’** (79%) compared to **AA’** and **CC’** (98% and 99%, respectively; **Figure S16A, Table S4**). Additionally, the two interhelical E@*g*-N@*a* contacts (E@*g*’-N@*a* / E@*g*-N@*a*’) that surround the Asn-Asn’ bond are engaged in hydrogen bonding for 64%/85% and 89%/66% of the simulation time for **AA’** and **CC’** (**Figure S16B**,**C** and **Table S5**). This long-lived interaction is absent for **BB’** as Lys precedes Asn for **B** and **B’** (**Table 1**). The presence of interhelical E@g-N@a hydrogen bonds corroborate a previous report by Hodges and co-workers, which disclosed the stabilizing effect of this particular interaction within the crystal structure of cortexillin I/GCN4 hybrid CC peptides.^42^ They revealed that E@g-N@a interactions form a network of hydrogen bonds that stabilize the Asn-Asn’ core region of the dimer.

Snapshots of the MD simulations for **AA’** and **CC’** reveal a slightly modified version of the previously reported hydrogen bonding network.^42^ In our case, using MD simulations, we observe a more extended hydrogen bonding network that connects across the *g*-*a*’-*a*-*g*’ layer (Glu-Asn’-Asn-Glu’; **Figure 4**). Within this network, peripheral E@*g* and E@*g*’ residues, which flank the interhelical Asn interaction, appear to “lock” the Asn pair in place via hydrogen bonds.

**Figure 4.**
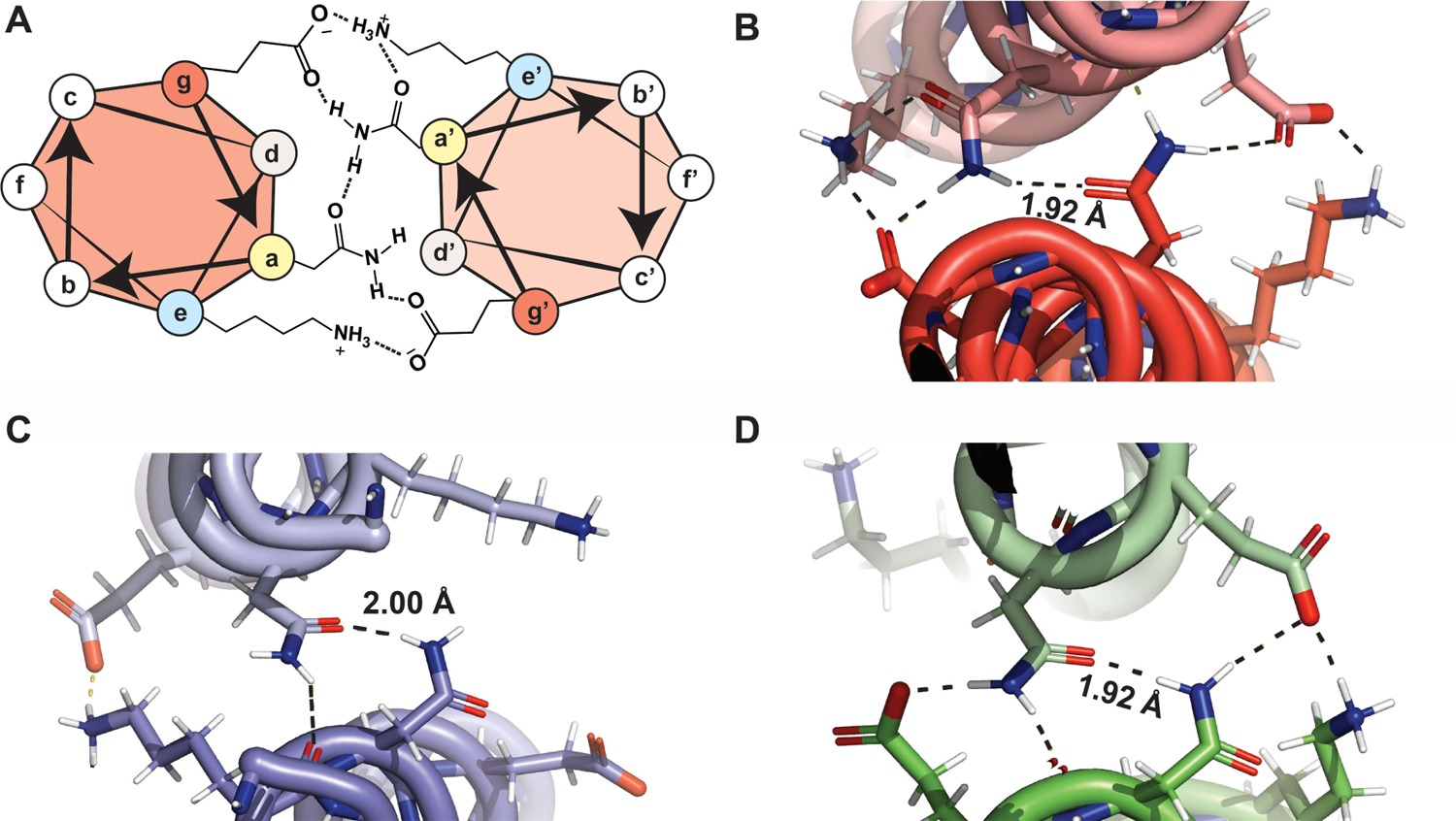
(A) Helical wheel diagram depicting the interhelical *g*-*a*’-*a*-*g*’ hydrogen bonding network that is observed when E@*g*-N@*a* contacts are present, including salt bridge contacts between *e* and *g* positions and a single intrahelical K@*e*’-N@*a*’ hydrogen bond. Snapshot of the MD simulations highlighting the hydrogen bonding interactions (dashed lines) that are present in proximity to the interhelical Asn-Asn’ contact for (**B**) **AA’**, (**C**) **BB’**, and (**D**) **CC’**. Average Asn-Asn’ bond distances are also displayed.

Additionally, MD trajectories show that Lys at *e* and *e’* readily form interchain salt bridges with the flanking Glu residues to create a more expansive hydrogen bonding network that spans across 6 residues (Lys’-Glu-Asn’-Asn-Glu’-Lys), and explains the increase in the number of salt bridges, on average, for **AA’** and **CC’** (**Figure S15**). The asymmetric engagement of E@*g*-N@*a*’ and E@*g*’-N@*a* interactions is attributed to the added stability associated with an intrahelical Lys-Asn hydrogen bond that can only form on one side of the CC interface (**Figure 4A**).^42^ From PDBePISA analysis, the prevalence of Glu-Lys contacts is markedly higher within the heptad that contains the Asn-Asn’ contact (**Figure S16D** and **Table S6**), indicating that the Glu-Asn’-Asn-Glu’ hydrogen bonding network spatially orients the E@*g*/*g*’ in an ideal configuration that is predisposed for forming interhelical salt bridges (**Figure 4A**). The presence of Lys at the *g* and *g*’ sites prevents the organization of this extended hydrogen bonding network for **BB’**. In this case, the alignment of side-chain carbonyl groups of Asn in one direction, which are co-planar when engaged, inhibits one carbonyl from accepting hydrogens from the side-chain amine of Lys (**Figure 4A**). Furthermore, the extended length of K@*g*/*g*’ cannot accommodate both salt bridge formation (with Glu located across the helix) and a hydrogen bond with Asn.

The increase in stability afforded by the hydrogen bonding network present within **AA’** and **CC’** are substantiated through analysis of the bond distance and conformational dynamics of the Asn-Asn’ interaction. Shorter Asn-Asn’ bond distances of **AA’** and **CC’** (1.92 ± 0.15 Å and 1.92 ± 0.14 Å, respectively) compared to **BB’** (2.00 ± 0.49 Å) corroborate the tighter packing and hence increased stability of the former pairs (**Figure S17**).^45^ Previous studies have shown that the buried Asn side chains are highly dynamic, exchanging between three primary configurations that are based on whether the side-chain is directed towards the interface (“inside”, χ1 ≈ −60°, *gauche*), away from the interface (“out”, χ1 ≈ +60°, *gauche*^-^) or somewhere in-between (“middle”, χ1 ≈ ±180°, *trans*).^33^ Side-chain dihedral angles of all three on-target dimers consists of the preferred “inside-middle” (or identical “middle-inside”) arrangement (χ1 ≈ −70° and χ1 ≈ ±180°), which accommodates the hydrogen bonding between the interhelical Asn residues (**Figure S18**). However, although the dihedral angles remain unchanged for all three on-target pairs, a flip from “inside-middle” to “middle-inside” was only observed within the second simulation run for **BB’** (**Figure S18B,E**). While using much longer simulation times and/or employing alternative MD techniques with enhanced conformational sampling (*e.g.*, REMD) would be required to fully map the conformational space of interchain Asn interactions, at a significantly increased computational cost, our results support the hypothesis that the decreased thermostability of **BB’** is attributed to the weaker and more dynamic interchain Asn-Asn’ contact.

In summary, interhelical E@*g*-N@*a* hydrogen bonds, which have been previously reported,^42^ are responsible for coordinating a hydrogen bonding network that stabilizes the CC interface by “locking” the Asn-Asn’ contact in place. The presence of this bonding network is further stabilized through long-lived Glu-Lys salt bridges that form almost exclusively between Glu/Lys pairs that are in the immediate proximity to the Asn-Asn’ bond. Replacement of Glu with Lys at the proximal *g* and *g*’ position precludes the formation of this CC-stabilizing hydrogen bonding pattern, which leads to a weaker and more dynamic Asn-Asn’ interaction, and ultimately manifests via the lower CC thermostability of **BB’**. Moreover, our results underscore the ability of MD simulations to yield accurate structural and dynamical data quickly and relatively inexpensively compared to x-ray crystallography. In turn, this allows for rapid feedback between model-based design and experimental validation, which can then be leveraged to gain greater clarity in resolving the various competing interactions present within the CC interface.^33^

### Redesign of BB’ with Glu-Asn Interactions Increases CC Thermostability

We hypothesized that redesigning **BB’** with Glu preceding Asn should give rise to the hydrogen bonding network present within **AA’** and **CC’**, and thus enhance the stability of **BB’**. MD simulations were conducted on two redesigned **BB’** pairs having one (**B_EE_** and **B’_KK_**) or two (**B_EK_** and **B’_EK_**) Glu residues installed at the *g*/*g*’ positions within the *g*-*a*’-*a*-*g*’ layer comprising the interchain Asn-Asn’ bond (subscripts of the redesigned sequences denote the electrostatic pattern within the Asn-containing heptad; **Table 1**). If they work as predicted, these new sequence designs will add an exciting new degree of freedom in specifying the stability of heterodimeric CCs as a function of the number of interhelical E@*g*-N@*a* interactions. Our MD simulations of these new structures show that the RMSD and RMSF values are comparable to the original on-target heterodimers (**Figures 3A,B**, and **S19A**,**B**, **S20A**,**B**), but as hypothesized, the average number of hydrogen bonds for **B_EE_B’_KK_** and **B_EK_B’_EK_** are increased step-wise from ∼3.7 to ∼5.0, respectively, with the latter pairing commensurate with the average number of hydrogen bonds for **AA’** and **CC’ (Figure 3C**, **S19C, S20C**). Furthermore, the average Asn-Asn’ bond distance was shortened to 1.94 ± 0.16 Å and 1.91 ± 0.13 Å for **B_EE_B’_KK_** and **B_EK_B’_EK_**, respectively, suggesting that **B_EK_B’_EK_** is potentially more stable than the other on-target dimers (**Figure S19F**, **S20F**).

PDBePISA analysis reveals that only a single installation of interhelical E@*g*-N@*a* interactions (**B_EE_B’_KK_**) is required to bring the percent occurrence of the Asn-Asn’ contact on par with **AA’** and **CC’** (**Figure S16A, Table S4**). Furthermore, the predominant Glu-Lys salt bridge interaction is restored back to the Glu/Lys pairs that are peripheral to the Asn-Asn’ bond – a significant departure from **BB’**, in which the Glu-Lys contacts are most prevalent at the C-terminus (**Figure S16D**). Visualization of the MD trajectory for **B_EK_B’_EK_** confirms the full extension of the hydrogen bonding network (Lys’-Glu-Asn’-Asn-Glu’-Lys) that was present in **AA’** and **CC’**, but absent in **BB’** (**Figure 5B**). As expected, with only a single interhelical E@*g*-N@*a* contact available, only half of the hydrogen bonding network is observed for **B_EE_B’_KK_** (**Figure 5A**).

**Figure 5.**
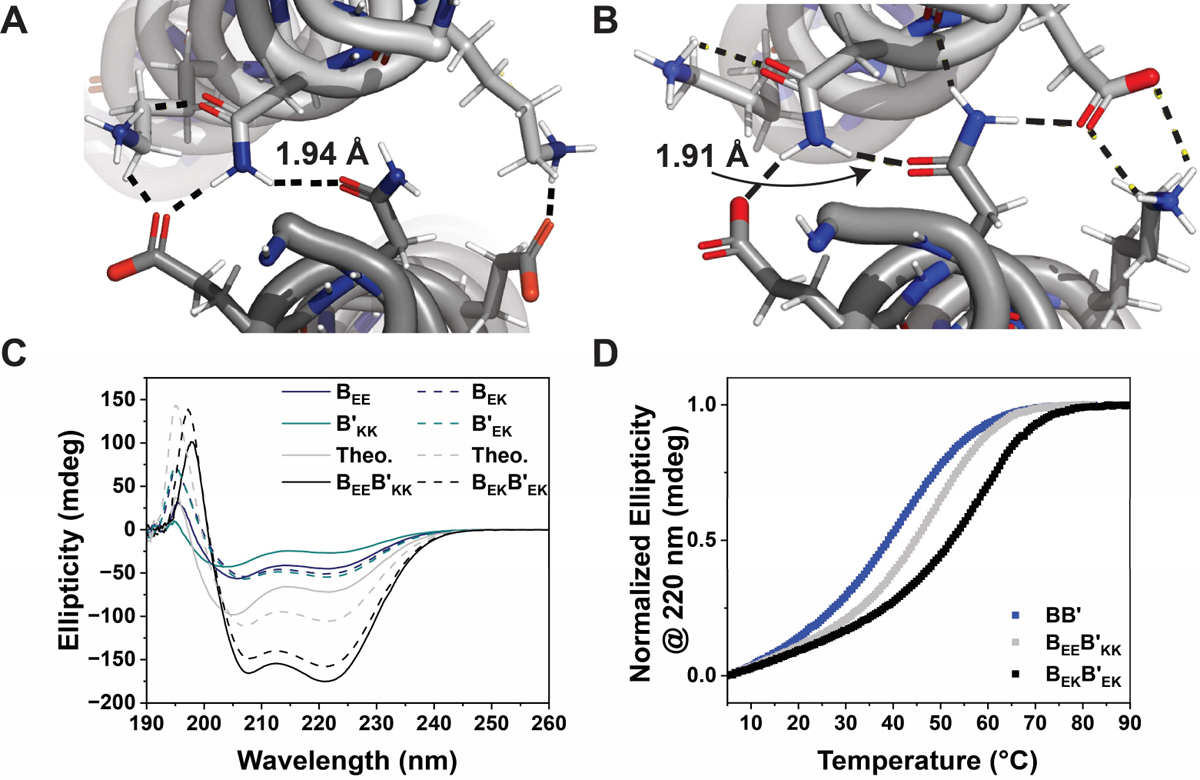
Snapshots of the MD simulation of (**A**) **B_EE_B’_KK_** and (**B**) **B_EK_B’_EK_** revealing the limited and extensive hydrogen bonding network, respectively, around the shortened interchain Asn-Asn’ bond. (**C**) CD spectra of **B_EE_**, **B’_KK_**, **B_EK_**, **B’_EK_**, **B_EE_B’_KK_**, and **B_EK_B’_EK_**. (**D**) Normalized thermal denaturation profiles of **BB’**, **B_EE_B’_KK_**, and **B_EK_B’_EK_**, which display the increase in T_m_ as a function of the number of interchain E@*g*-N@*a* contacts present within the dimer.

To test these theoretical predictions, the four new **B** variants were synthesized, isolated, and characterized (**Table 1**, **Figures S21-22**). The predictions from MD simulations of the redesigned sequences were successfully verified experimentally. CD and SEC data of the redesigned dimer species verify that they form dimeric CCs (**Figure 5C, S23**), and as expected, thermal denaturation of **B_EE_B’_KK_** and **B_EK_B’_EK_** reveal a step-wise increase in thermostability (T_m_ of 45°C and 53°C, respectively; **Figure 5D, S24**) in comparison to **BB’** (T_m_ = 39°C; **Figure 2E**). The T_m_ observed for **B_EK_B’_EK_** actually exceeds that of **AA’** and **CC’**. These results are consistent with prior studies that demonstrated that buried polar Asn-Asn’ interactions are less disruptive to the hydrophobic interface when excluded from the core region, and as a result, sequences with Asn incorporated near the termini exhibit greater thermostability.^30^ Thermodynamic parameters for **B_EE_B’_KK_** and **B_EK_B’_EK_** reveal the step-wise decrease in the free energy as a function of installing either one or two interhelical E@*g*-N@*a* contacts, respectively (**Table 2, Figure S25**). From this analysis, an individual Glu-Asn contact results in an approximate 3.48 kJ/mol increase in stability, when compared to **BB’**, while two Glu-Asn contacts result in a 5.53 kJ/mol increase in stability. These results are consistent with the prior work that reported a 0.76 kcal/mol (3.2 kJ/mol) increase in stability per interchain E@*g*-N@*a* pair.^42^ Lastly, we note a 4x and 10x decrease in K_d_ values for **B_EE_B’_KK_** and **B_EK_B’_EK_**, respectively (**Table 2**). Altogether, the results demonstrate that the CC stability within our *de novo* peptide set can be systematically modulated by controlling the number of interhelical E@*g*-N@*a* interactions.

## 4. Conclusion

We establish a *de novo* set of orthogonal CC peptides that comprise 3.5 heptads in length and contains a single buried polar interaction to dictate dimer specificity. CD analysis of the on- and off-target heterodimers confirm the fidelity of the designed on-target pairs, and is consistent with results from MD simulations, which identify differential interactions between on- and off-target dimers, thus providing a straightforward method for predicting (and subsequently adding) future orthogonal CCs into the collection. In addition, comprehensive analysis of the MD data uncovers two key design considerations that can be employed for tuning the thermostability and binding strength between similarly designed orthogonal CC sequences. First, interhelical Glu-Asn interactions coordinate a long-range E@*g*-N@*a*’-N@*a*-E@*g*’ hydrogen bonding network that stabilizes the CC interface by locking the interhelical Asn-Asn’ contact in place. This leads to a substantial ∼10-fold decrease in K_d_ (>14°C increase in T_m_) when compared to similar CC sequences that do not contain interhelical E@*g*-N@*a* bonds. This design consideration provides an additional handle to modulate the stability of commonly utilized Asn-containing CCs with minimal alterations to the peptide sequence design. Second, Glu-Lys salt bridging contacts, which are available throughout the CC, occur much more frequently for Glu/Lys pairs that immediately flank the Asn-Asn’ bond, and contribute to the organization of a more expansive 6-residue hydrogen bonding network (Lys’-Glu-Asn’-Asn-Glu’-Lys) that provides additional stability around the Asn-Asn’ contact. We envision that this second consideration will help direct future orthogonal CC sequence designs by focusing positive and negative design elements within the heptad(s) that comprise(s) Asn, leading to increased energy gap differences between on- and off-target designs. In conclusion, our results highlight the utility of MD simulation to contribute additional CC sequence design rules that can be incorporated into the CC toolkit and allow for the construction of more complex and dynamic architectures with greater level of control over their properties.

## Acknowledgements

This work was supported by the University of California, Merced. A.R.P. acknowledges fellowship support from the NSF-CREST: Center for Cellular and Biomolecular Machines at UC Merced (NSF-HRD-1547848 and NSF-HRD-2112675). Y.L. acknowledges fellowship support from UC LEADS (funded by the University of California Office of the President) and LAEP (funded by the California Student Aid Commission). We acknowledge computing time on the Pinnacles cluster at UC Merced (NSF-MRI-2019144). We thank Prof. Son Nguyen for use of the DLS instrument and Prof. Shahar Sukenik for helpful comments and discussion. We also thank Dr. Lauren Stark for access to her PDBePISA driver and analysis scripts, and Joseph McTiernan for help in writing our MDAnalysis scripts.

## TOC Figure

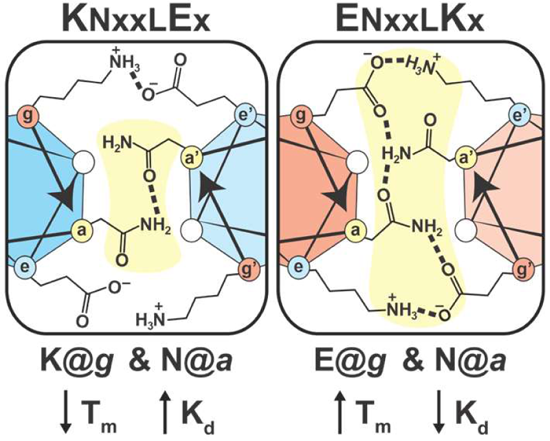

