## Supporting Information for "Interhelical E@*g*-N@*a* Interactions Modulate Coiled Coil Stability within a *De Novo* Set of Orthogonal Peptide Heterodimers"

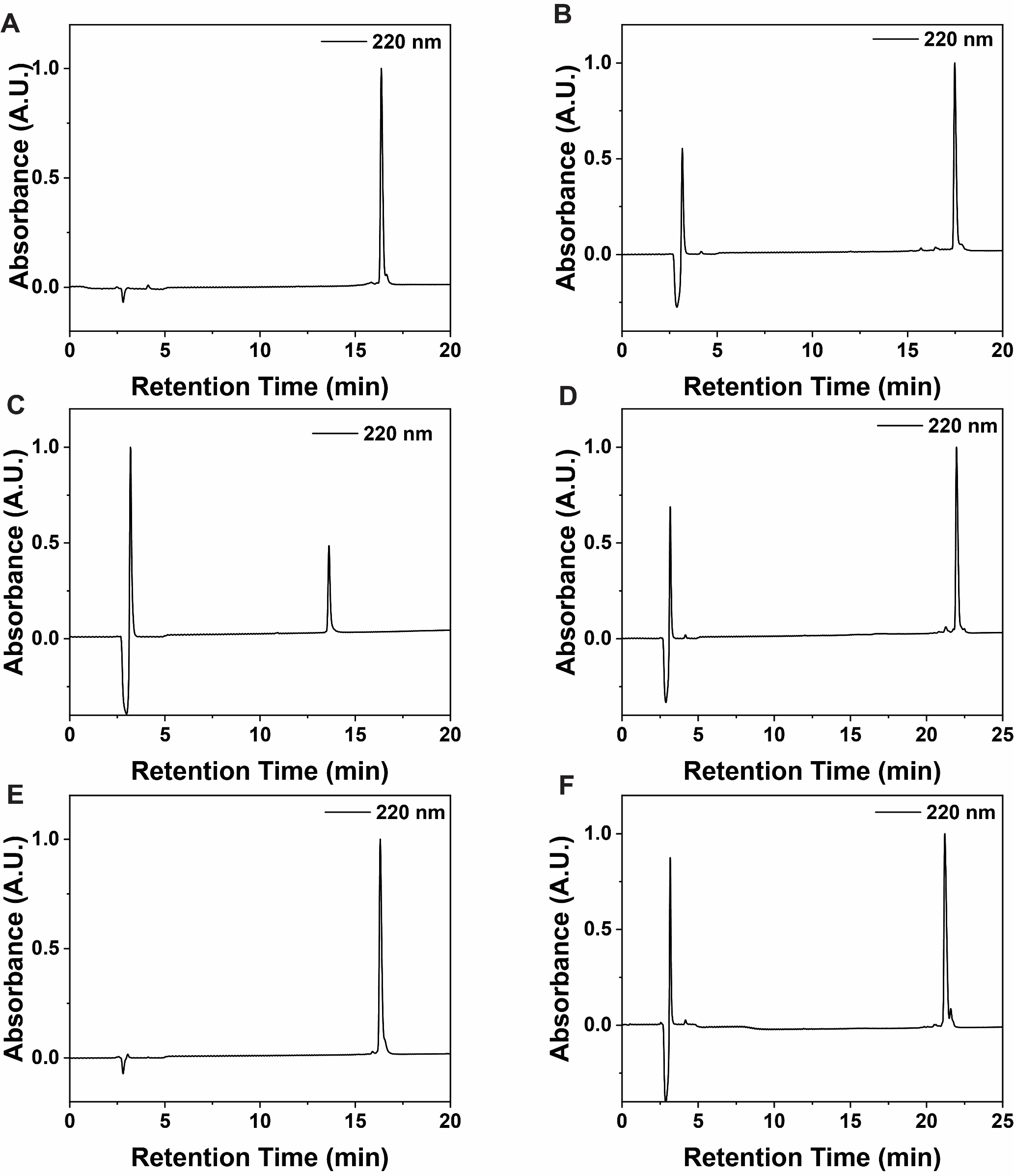

**Figure S1.** HPLC traces of (**A**) pep **A**, (**B**) pep **A’**, (**C**) pep **B**, (**D**) pep **B’**, (**E**) pep **C**, (**F**) pep **C’**.

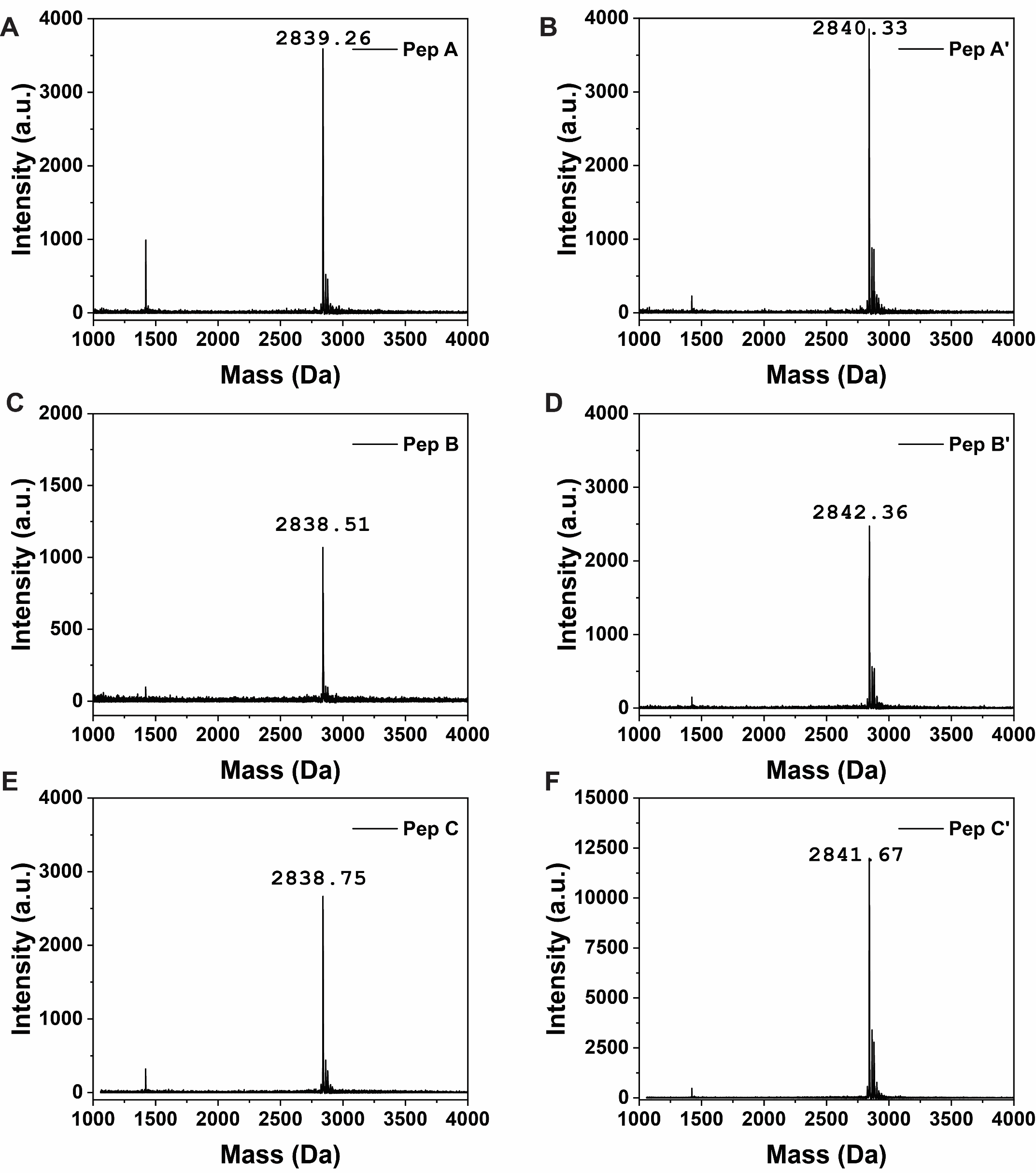

**Figure S2.** MALDI-TOF MS spectra of purified (**A**) **A**, m/z = 2839.26 [M+H]^+^, (**B**) **A’**, m/z = 2840.33 [M+H]^+^, (**C**) **B**, m/z = 2838.51 [M+H]^+^, (**D**) **B’**, m/z = 2842.36 [M+H]^+^, (**E**) **C**, m/z = 2838.75 [M+H]^+^, (**F**) **C’**, m/z = 2841.67 [M+H]^+^.

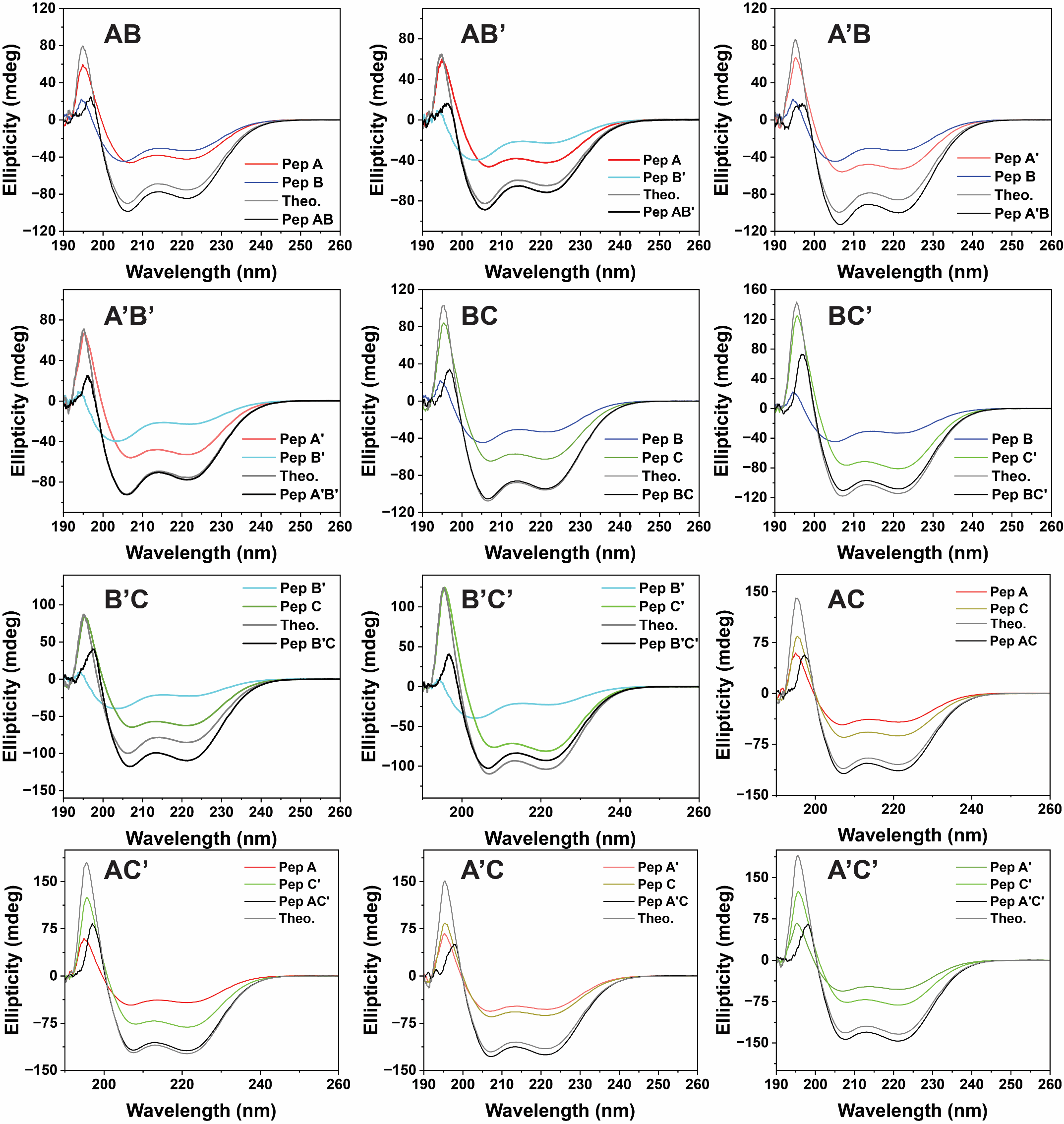

**Figure S3.** CD spectra of off-target CC dimers.

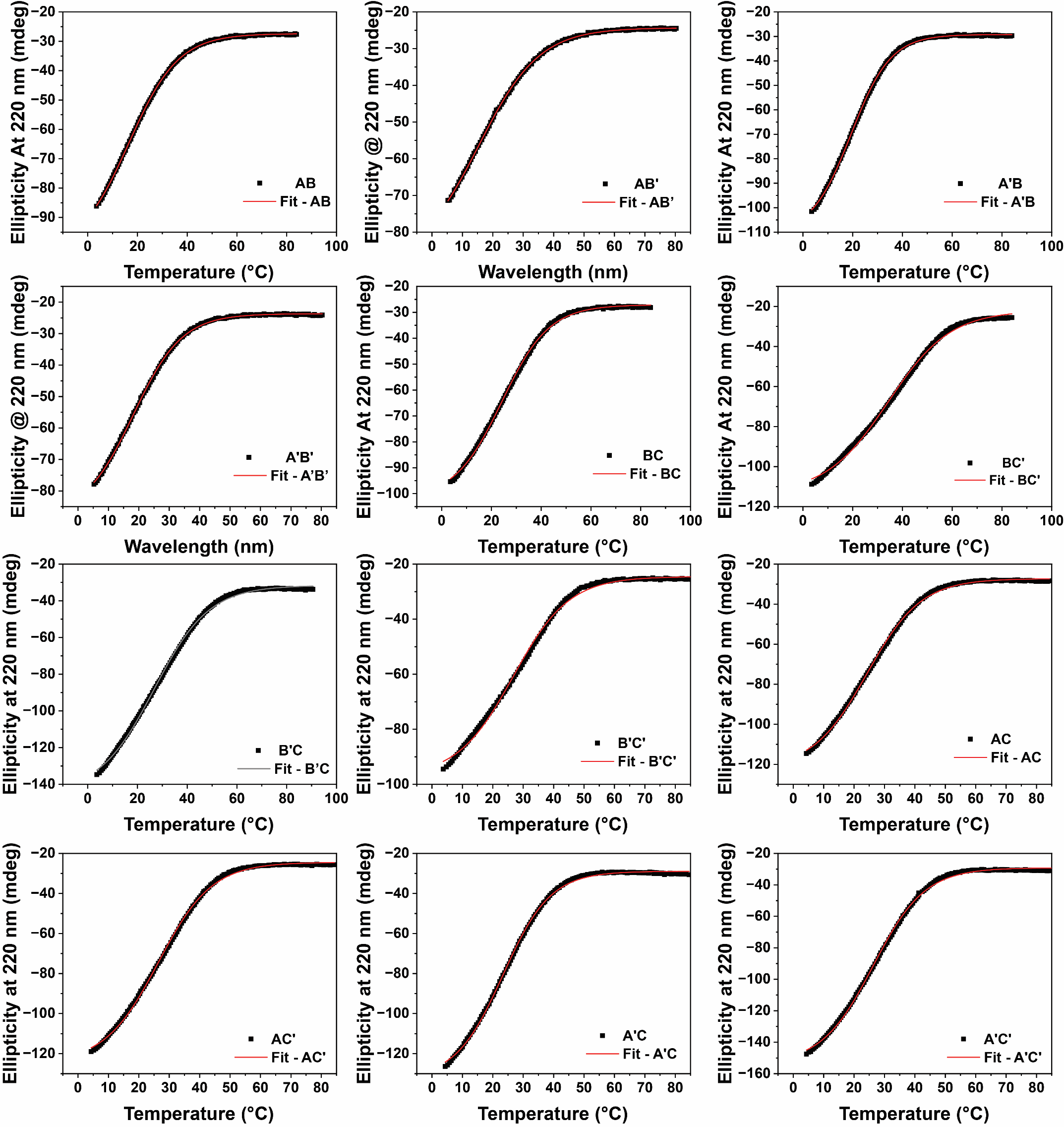

**Figure S4.** CD thermal denaturation plots and their fit curve for the twelve off-target interactions.

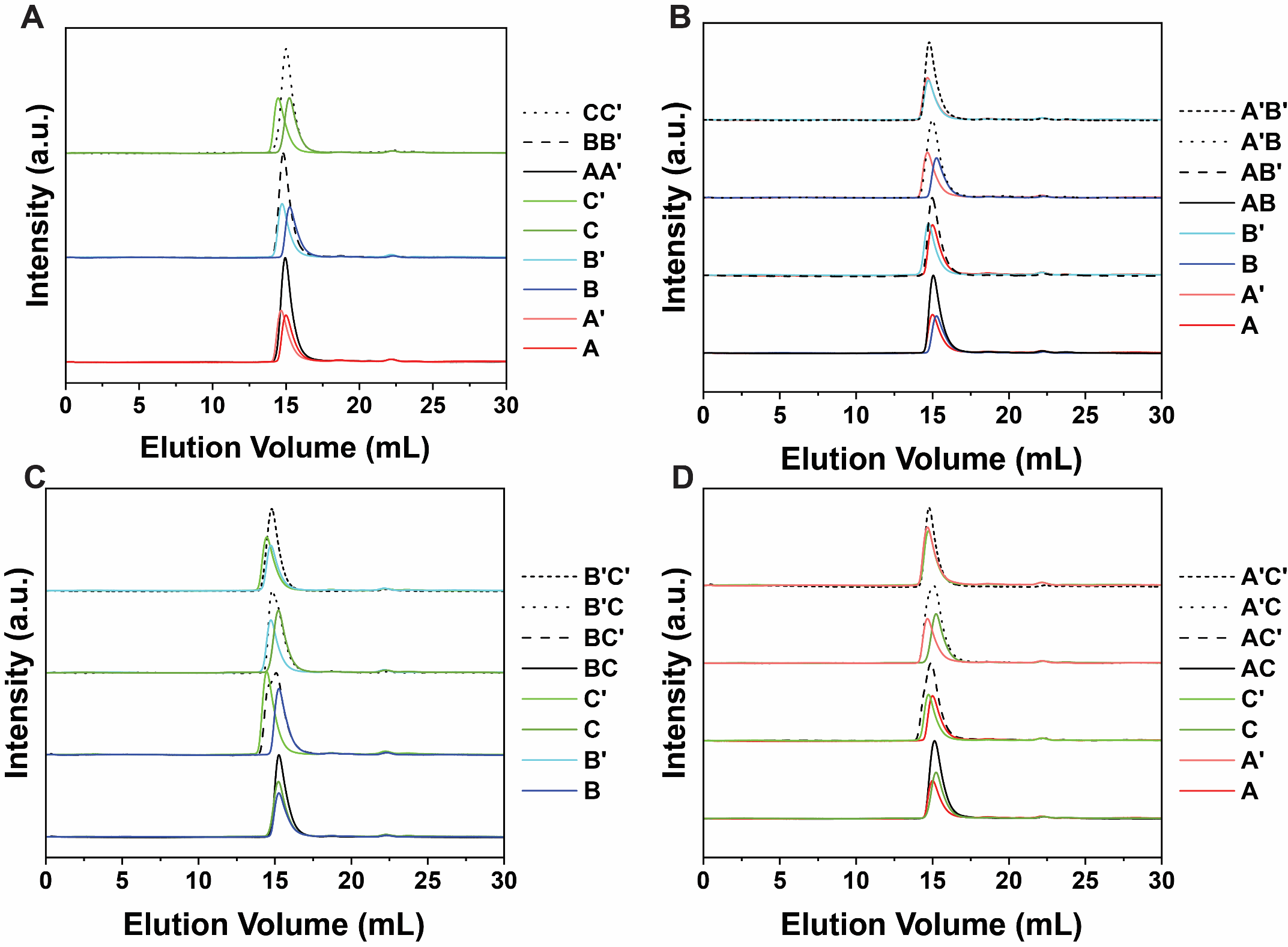

**Figure S5.** SEC profiles for (**A**) on-target interactions (**AA’**, **BB’**, and **CC’**) and monomers, (**B**) A/B peptide series, (**C**) B/C peptide series, and (**D**) A/C peptides series.

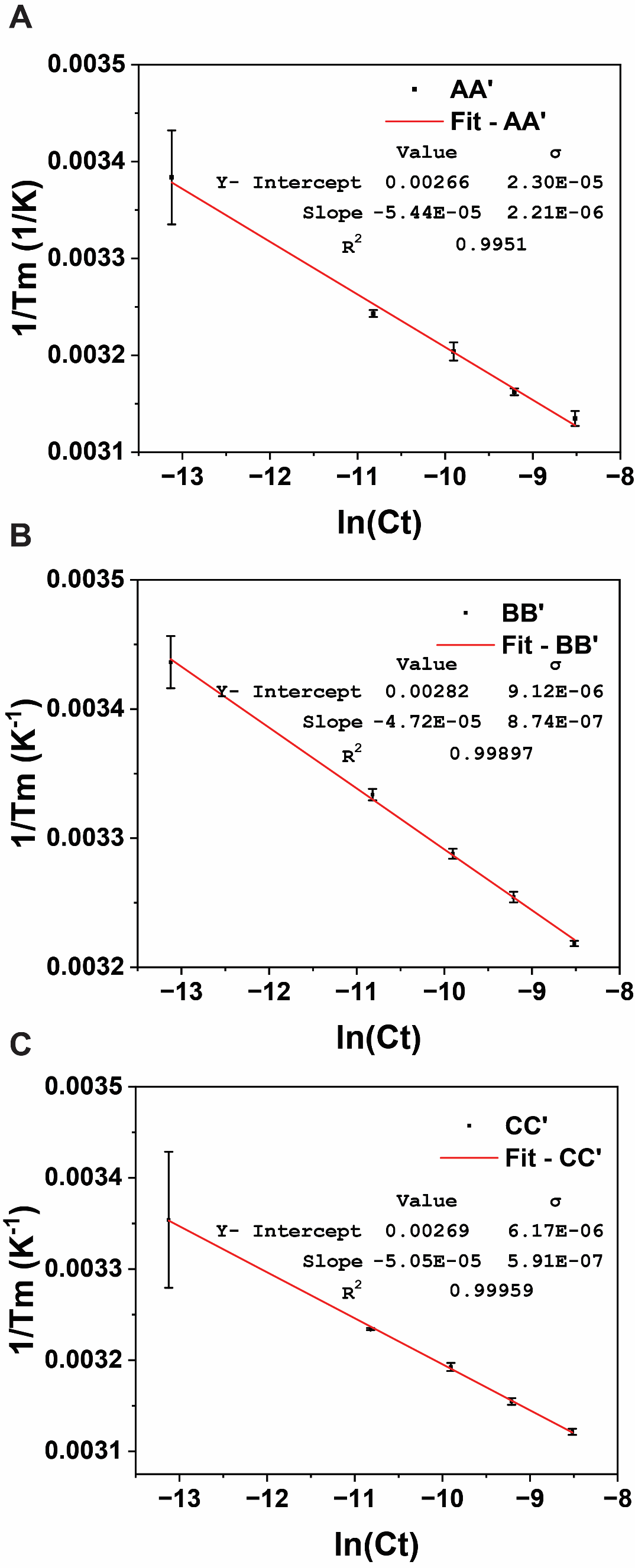

**Figure S6.** Van’t Hoff plots for the three on-target dimers: (**A**) **AA’**, (**B**) **BB’**, and (**C**) **CC’**.

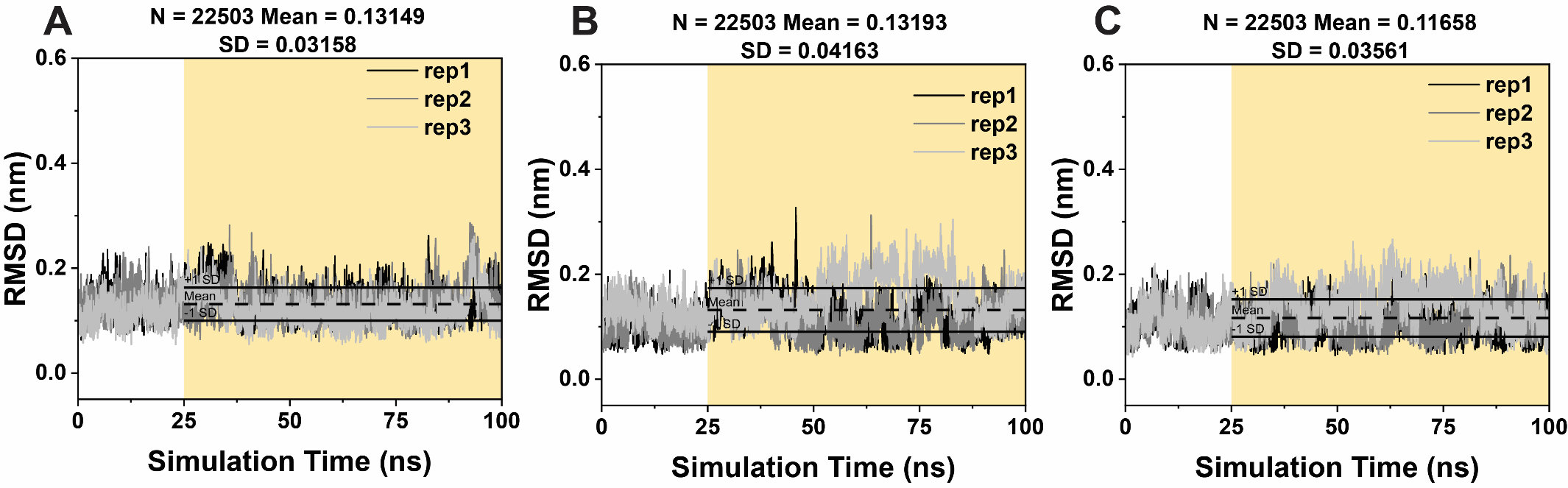

**Figure S7.** RMSD plots and analysis of on-target CC heterodimers: (**A**) **AA’**, (**B**) **BB**’, and (**C**) **CC’**. Mean and standard deviation values were calculated from the last 75 ns of simulation (shaded yellow portion).

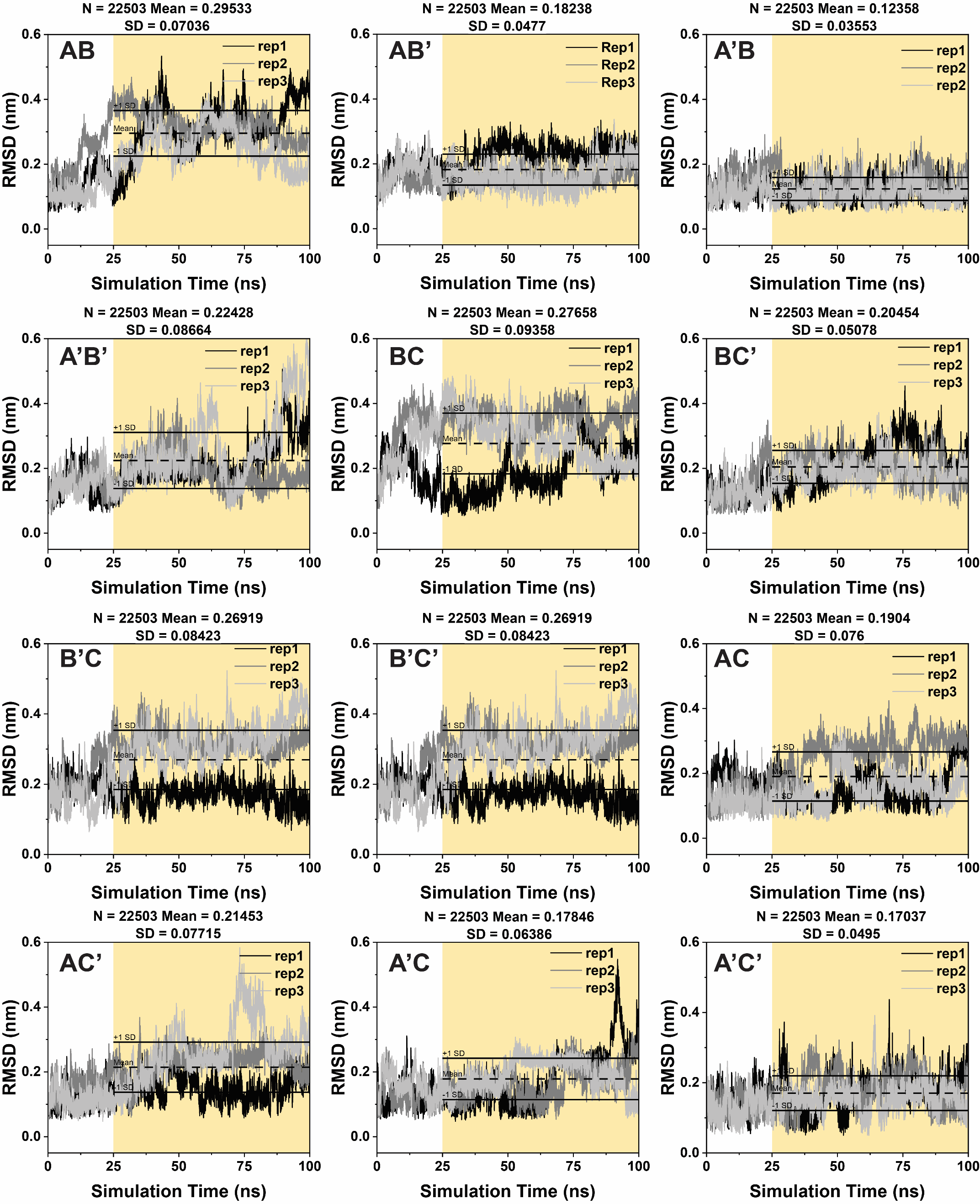

**Figure S8.** RMSD plots for the twelve off-target interactions. Mean and standard deviation values were calculated from the last 75 ns of simulation (shaded yellow portion).

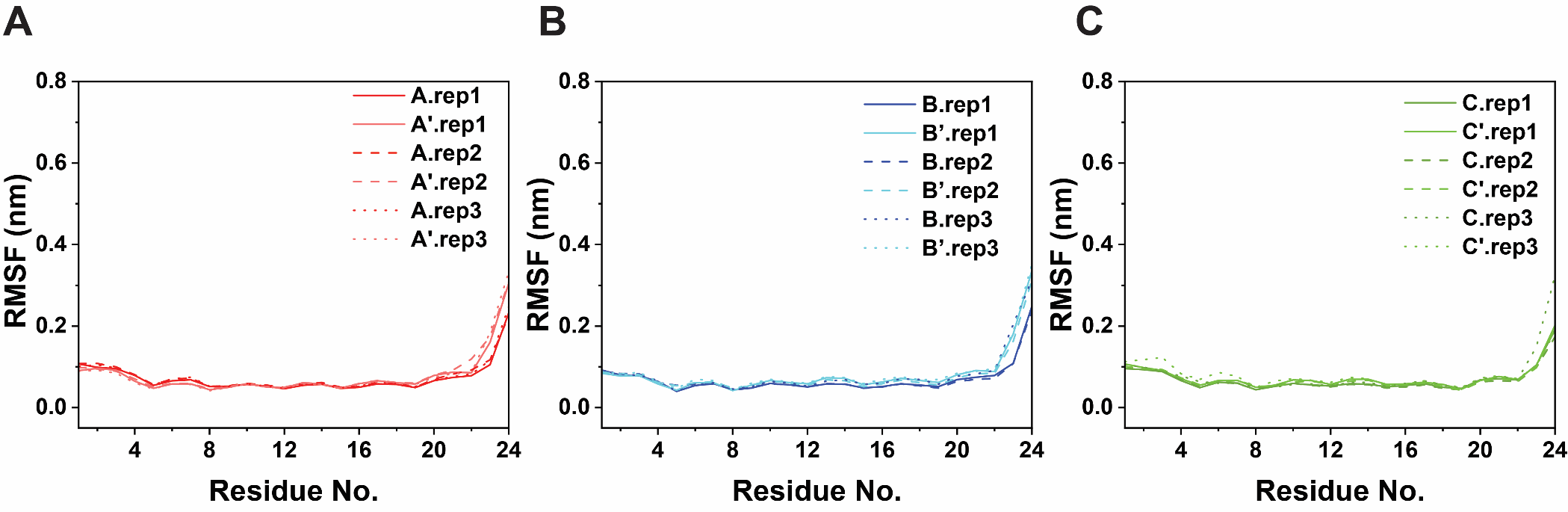

**Figure S9.** RMSF plots of on-target CC heterodimers: (**A**) **AA’**, (**B**) **BB**’, and (**C**) **CC’**.

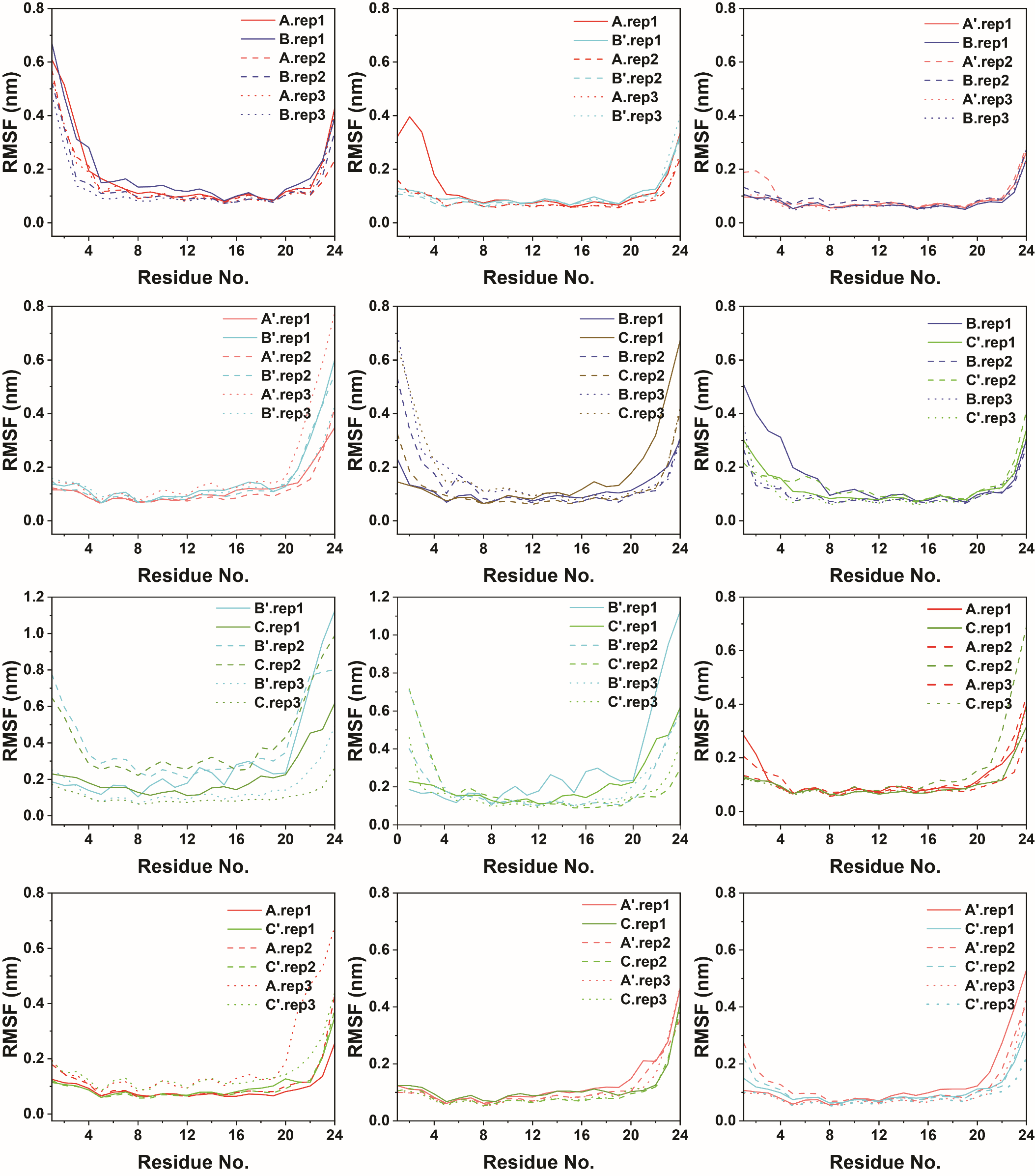

**Figure S10.** RMSF plots for the twelve off-target interactions.

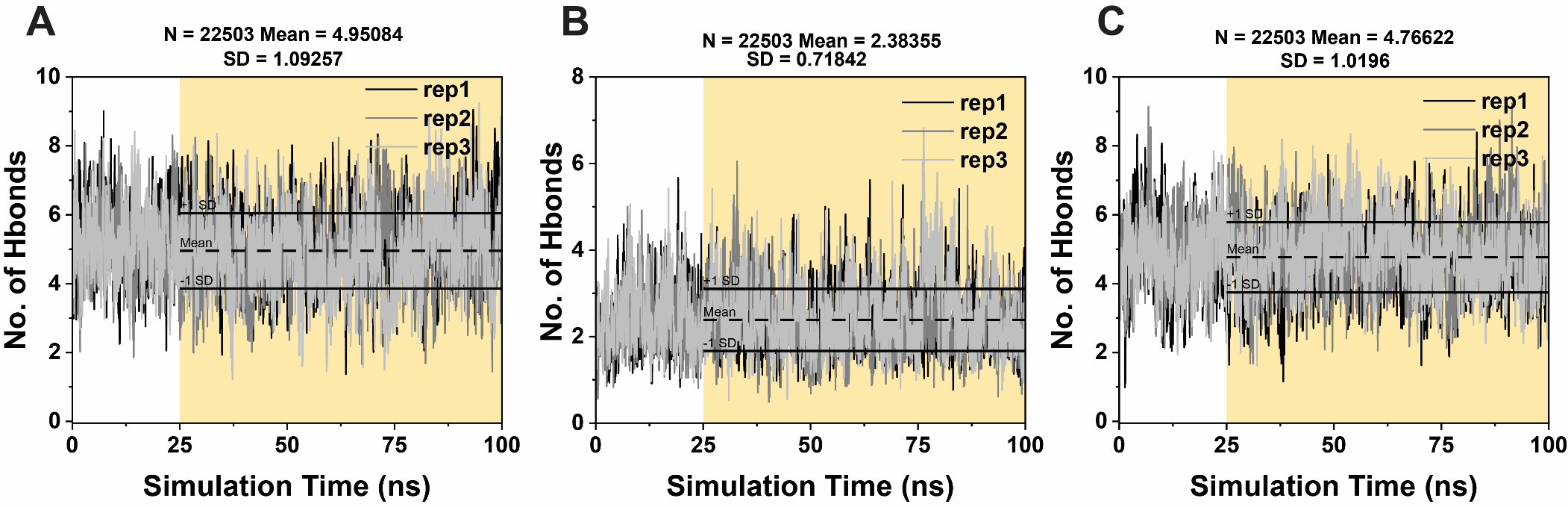

**Figure S11.** Number of interchain hydrogen bond contacts for each on-target dimer: (**A**) **AA’**, (**B**) **BB**’, and (**C**) **CC’**. Mean and standard deviation values were calculated from the last 75 ns of simulation (shaded yellow portion).

.

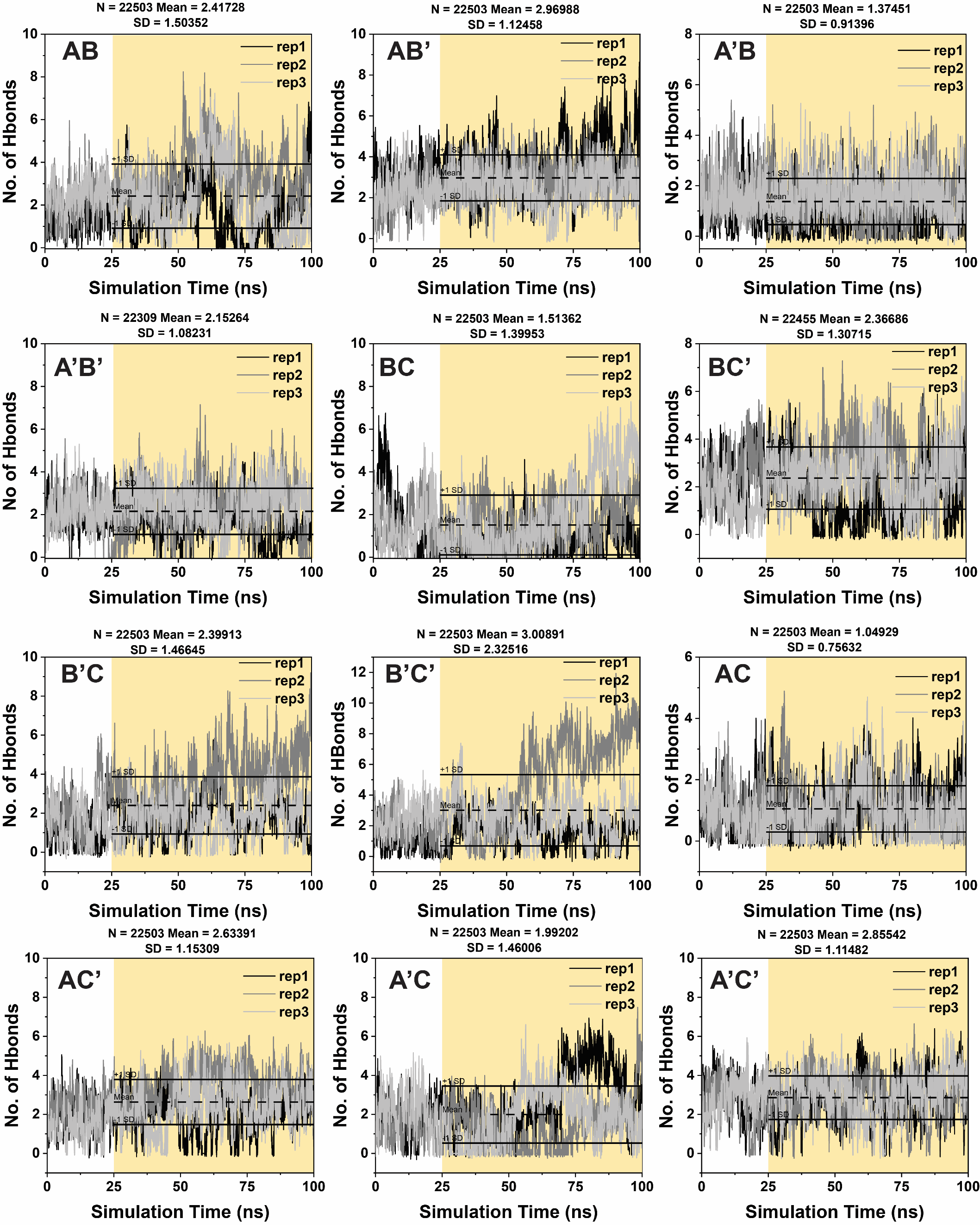

**Figure S12.** Number of interchain hydrogen bond contacts for the twelve off-target interactions. Mean and standard deviation values were calculated from the last 75 ns of simulation (shaded yellow portion).

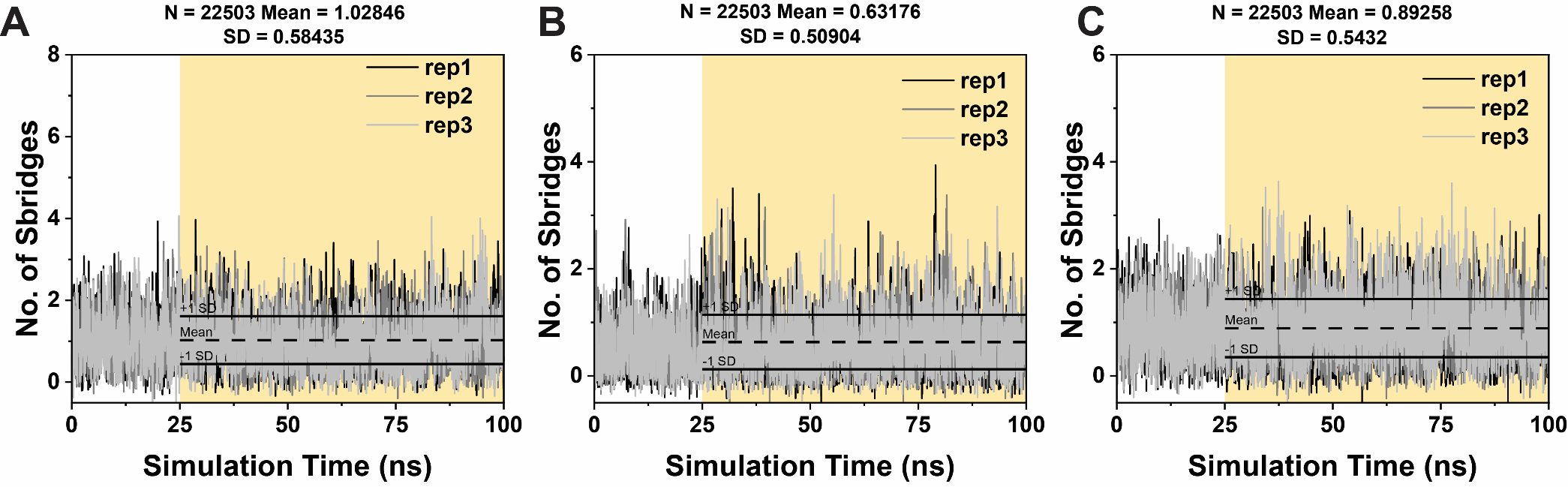

**Figure S13.** Number of interchain salt bridge contacts for each on-target dimer: (**A**) **AA’**, (**B**) **BB**’, and (**C**) **CC’**. Mean and standard deviation values were calculated from the last 75-ns of simulation (shaded yellow portion).

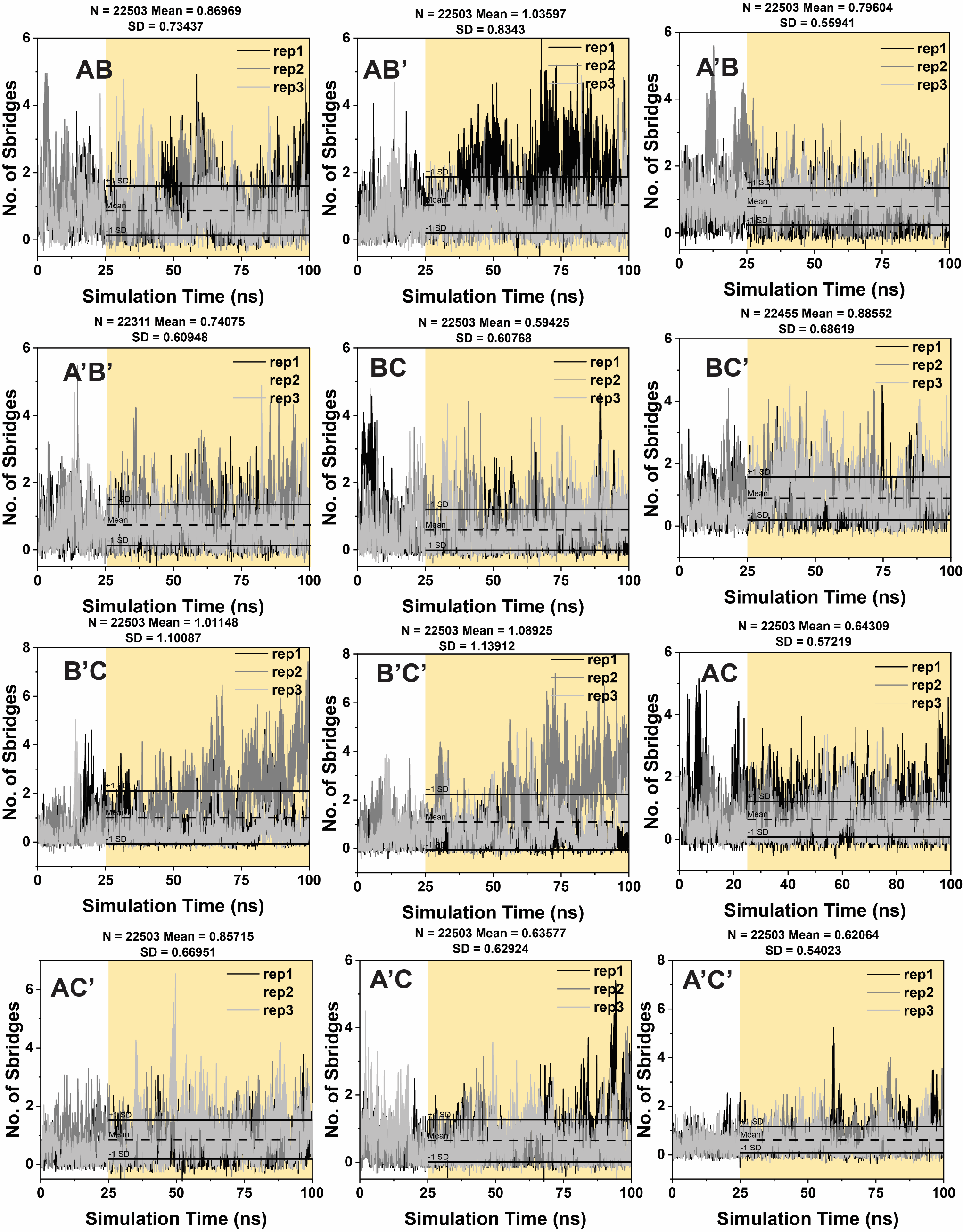

**Figure S14.** Number of interchain salt bridge contacts for the twelve off-target interactions. Mean and standard deviation values were calculated from the last 75 ns of simulation (shaded yellow portion).

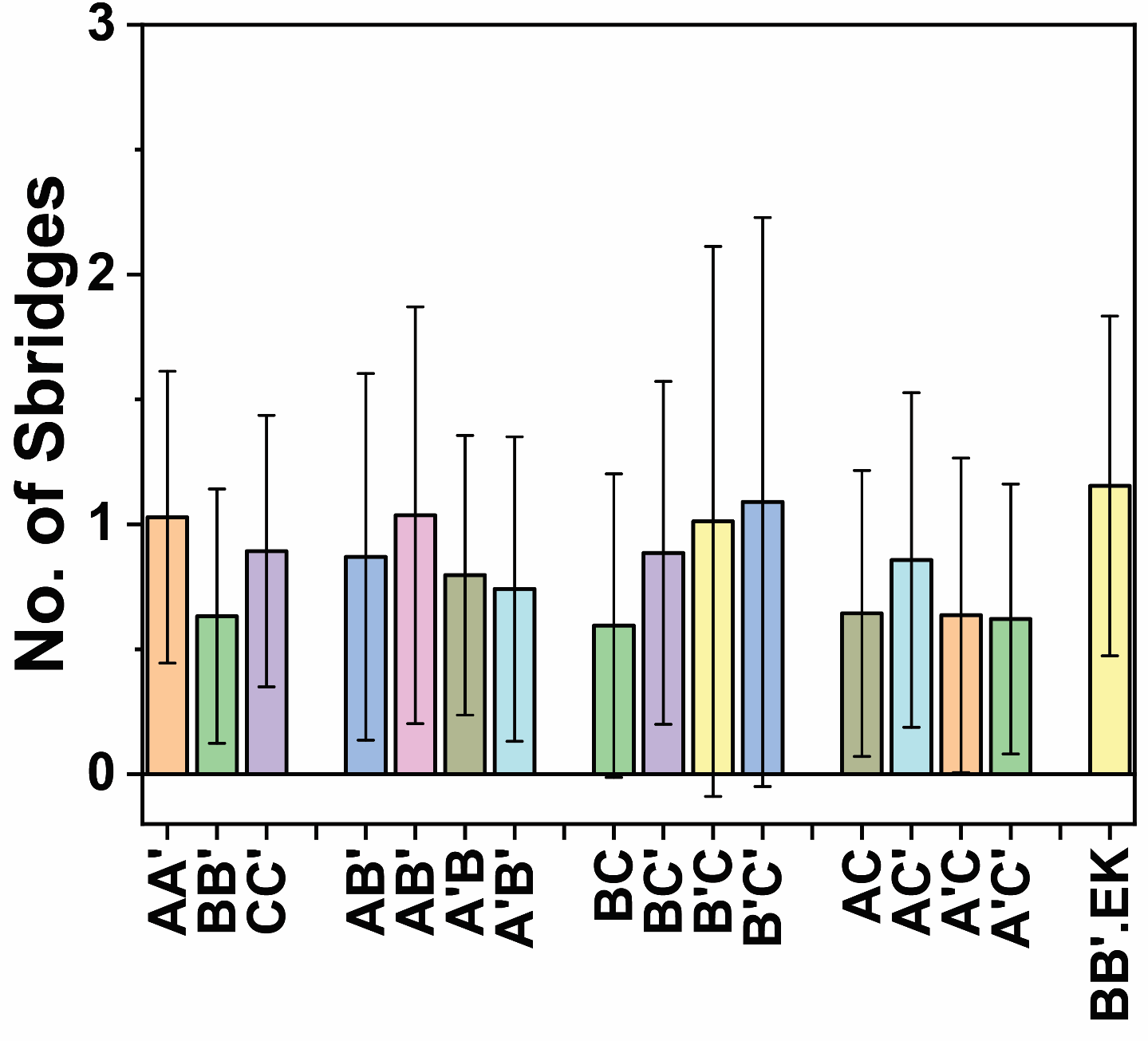

**Figure S15.** Average number of salt bridge contacts for all 15 on- and off-target pairings.

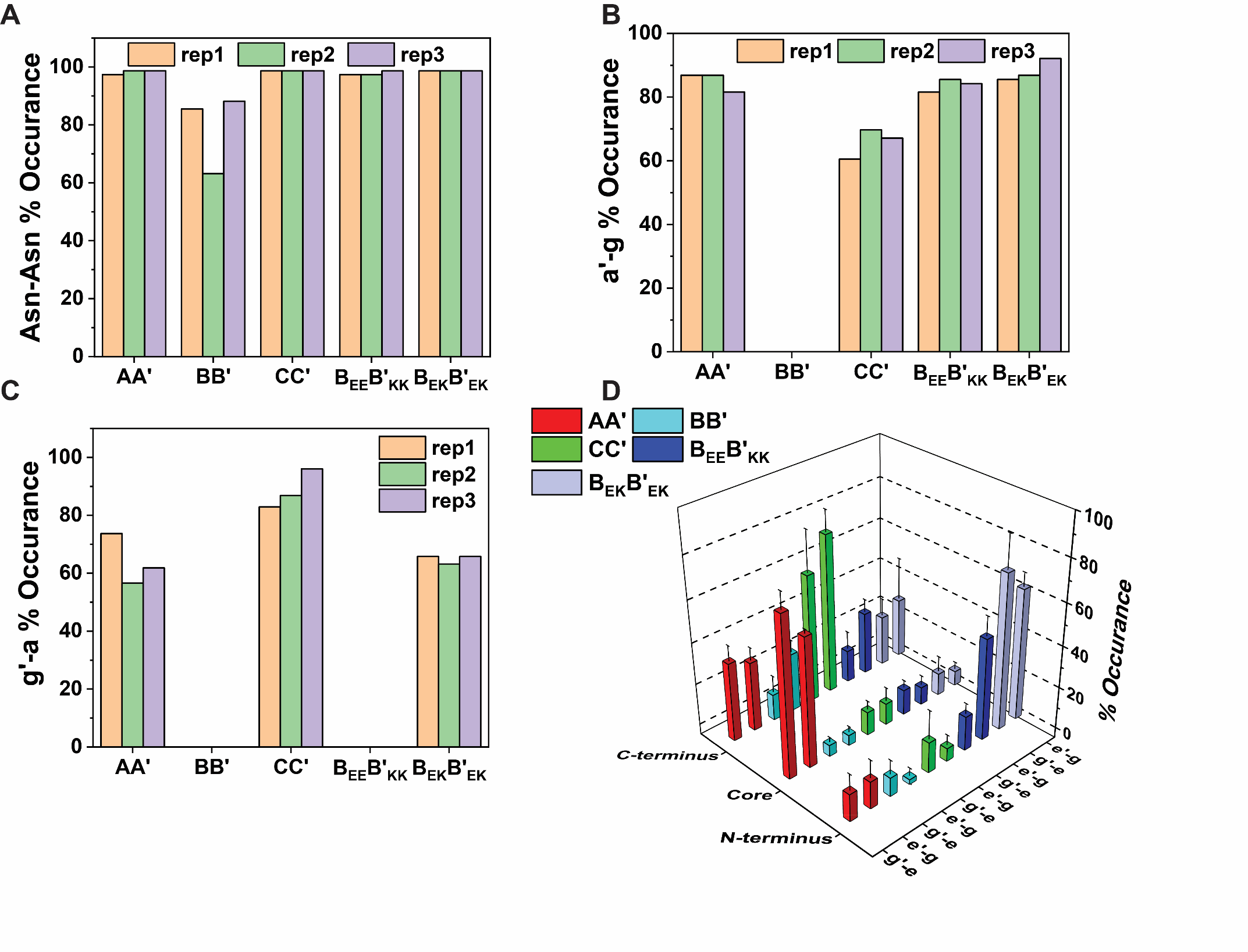

**Figure S16.**  PDBePISA analysis. (**A**) The percent occurrence of Asn-Asn’ contacts for the five designed dimers. (**B**, **C**) The percent occurrence of *a’-g­* and *g’-a* contacts (*i.e.* Asn’-Glu and Glu’-Asn). (**D**) The percent occurrence of salt bridges with respect to their locant in the peptide sequence, where C-terminus, Core, and N-terminus refer to the heptad at the C-terminus, core and N-terminus, respectively.

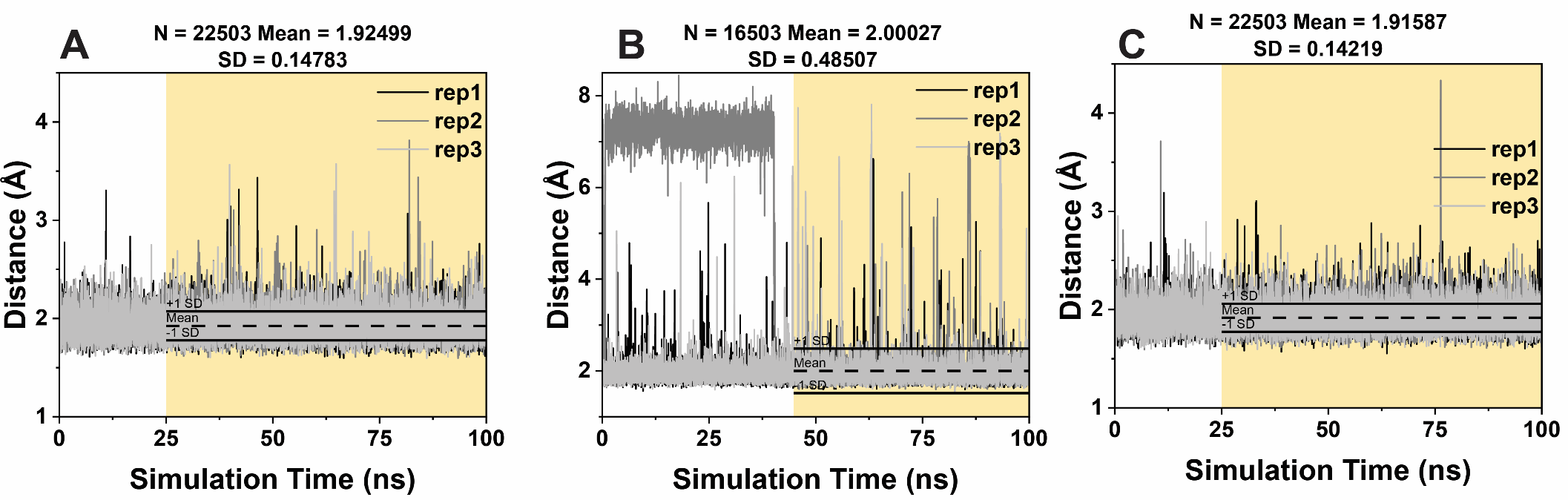

**Figure S17.** Asn-Asn’ hydrogen bond distance for (**A**) **AA’**, (**B**) **BB**’, and (**C**) **CC’**. Mean and standard deviation values were calculated from the last 75 ns of simulation, except for **BB’** due to conformational shift (shaded yellow portion).

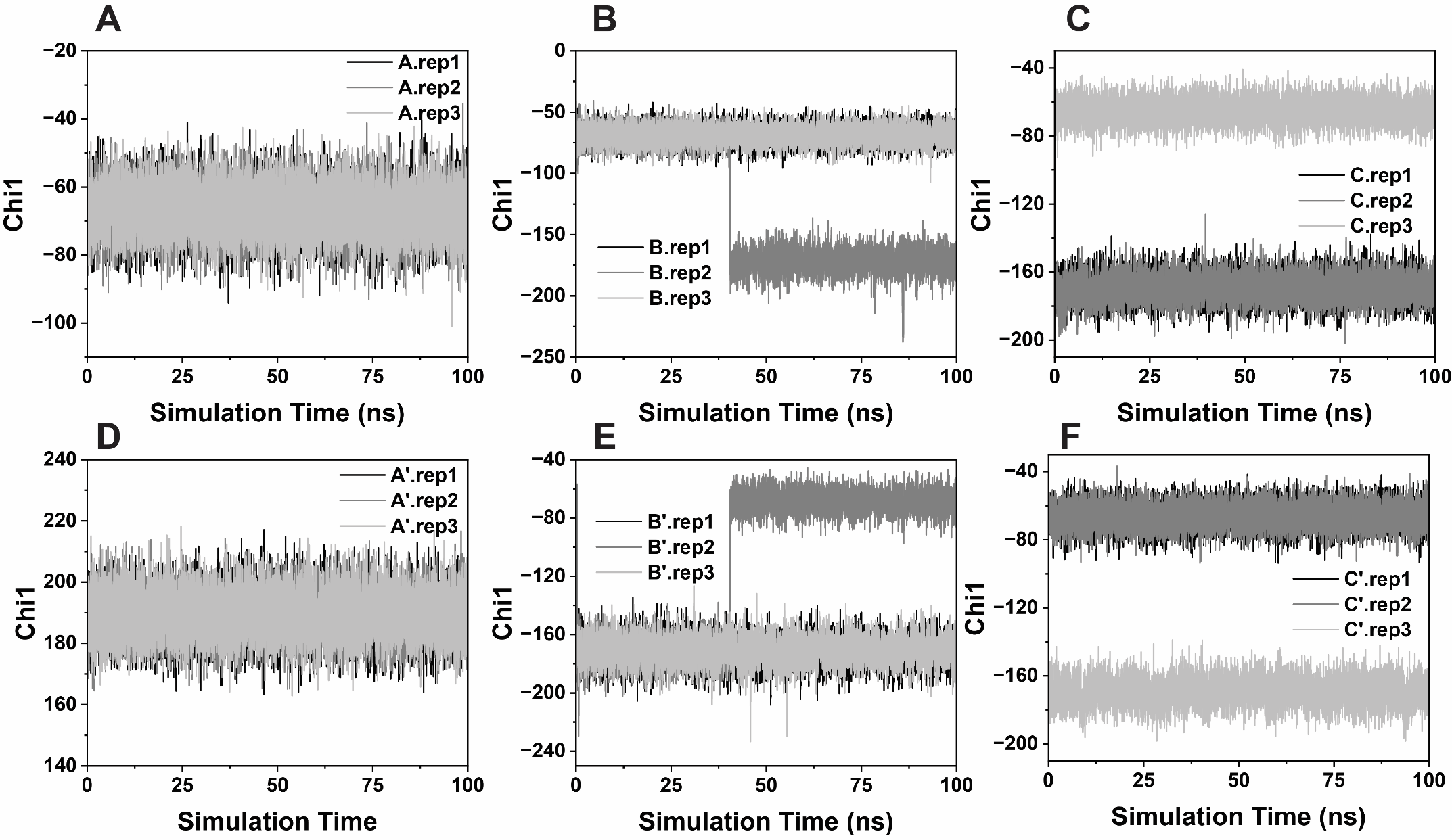

**Figure S18.** Dihedral angles of Asn side-chains of the on-target dimers: (**A**,**D**) **A** and **A’**, respectively, within **AA’**; (**B**,**E**) **B** and **B’**, respectively, within **BB’**; (**C**,**F**) **C** and **C’**, respectively, within **CC’**.

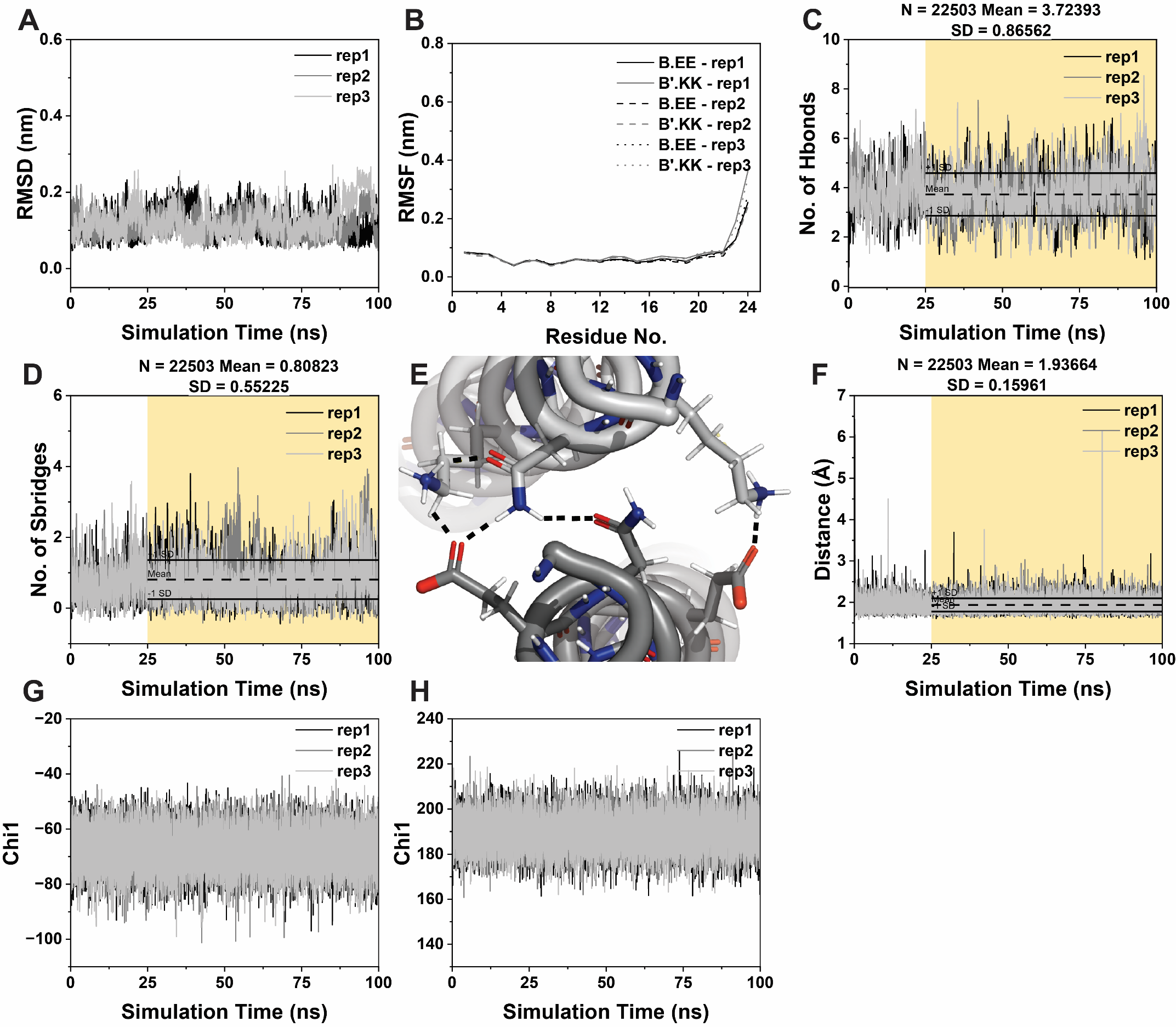

**Figure S19.** MD simulation results for **B_EE_B’_KK_**. Plots of (**A**) RMSD data, (**B**) RMSF data, (**C**) number of hydrogen bonds, and (**D**) number of salt bridge contacts. (**E**) Snapshot of a single MD trajectory highlighting the more limited hydrogen bonding network within **B_EE_B’_KK_**. (**F**) Asn-Asn’ hydrogen bond distance. Dihedral angles of Asn side-chain of (**G**) **B_EE_** and (**H**) **B’_KK_** within **B_EE_B’_KK_**. In panels **B** and **E**, dark gray is **B_EE_** and light gray is **B’_KK_**. Mean and standard deviation values were calculated from the last 75 ns of simulation (shaded yellow portion).

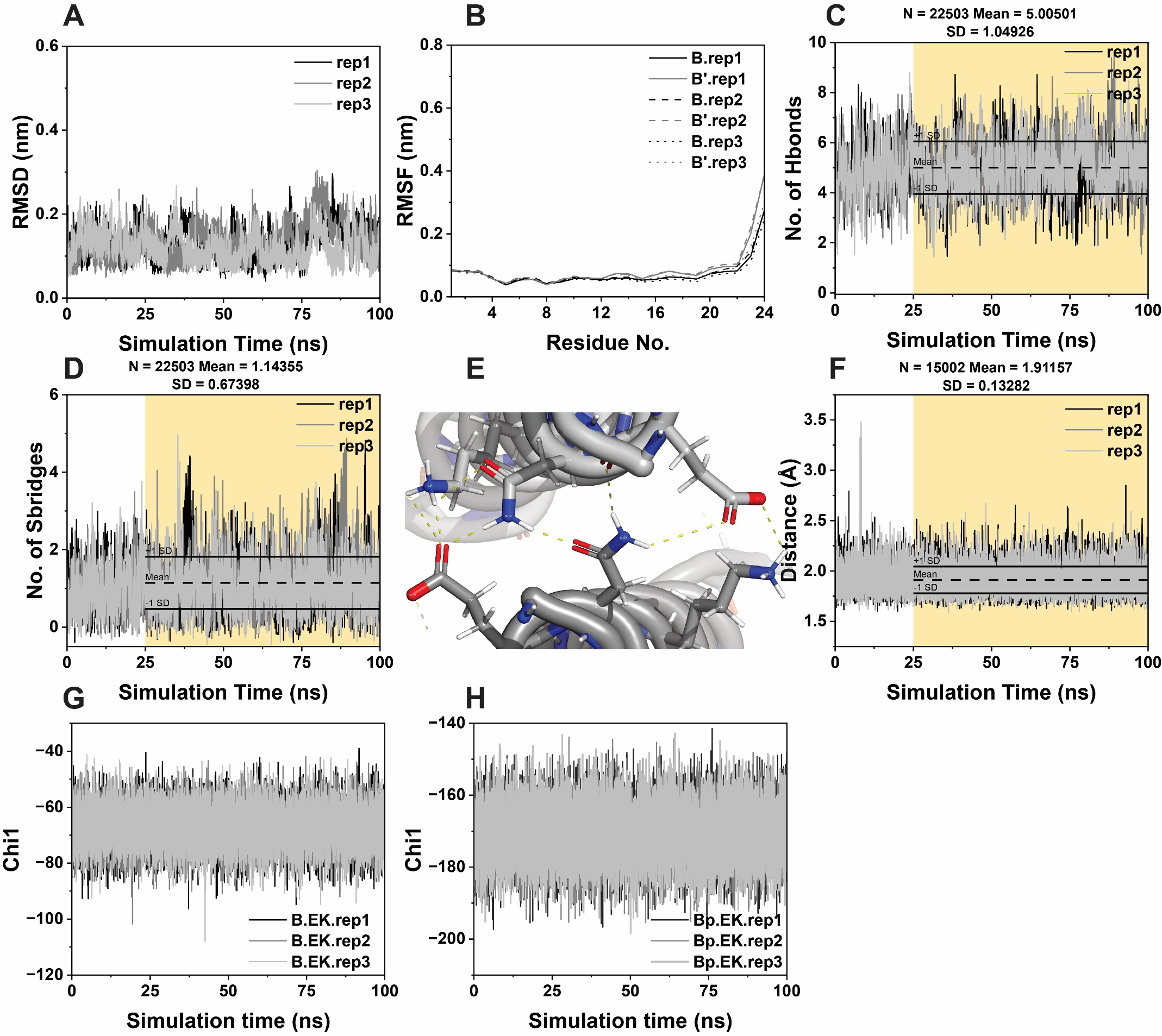

**Figure S20.** MD simulation results for **B_EK_B’_EK_**. Plots of (**A**) RMSD data, (**B**) RMSF data, (**C**) number of hydrogen bonds, and (**D**) number of salt bridge contacts. (**E**) Snapshot of a single MD trajectory highlighting the extensive hydrogen bonding network within **B_EK_B’_EK_**. (**F**) Asn-Asn’ hydrogen bond distance. Dihedral angles of Asn side-chain of (**G**) **B_EK_** and (**H**) **B’_EK_** within **B_EK_B’_EK_**. In panels **B** and **E**, dark gray is **B_EK_** and light gray is **B’_EK_**. Mean and standard deviation values were calculated from the last 75 ns of simulation (shaded yellow portion).

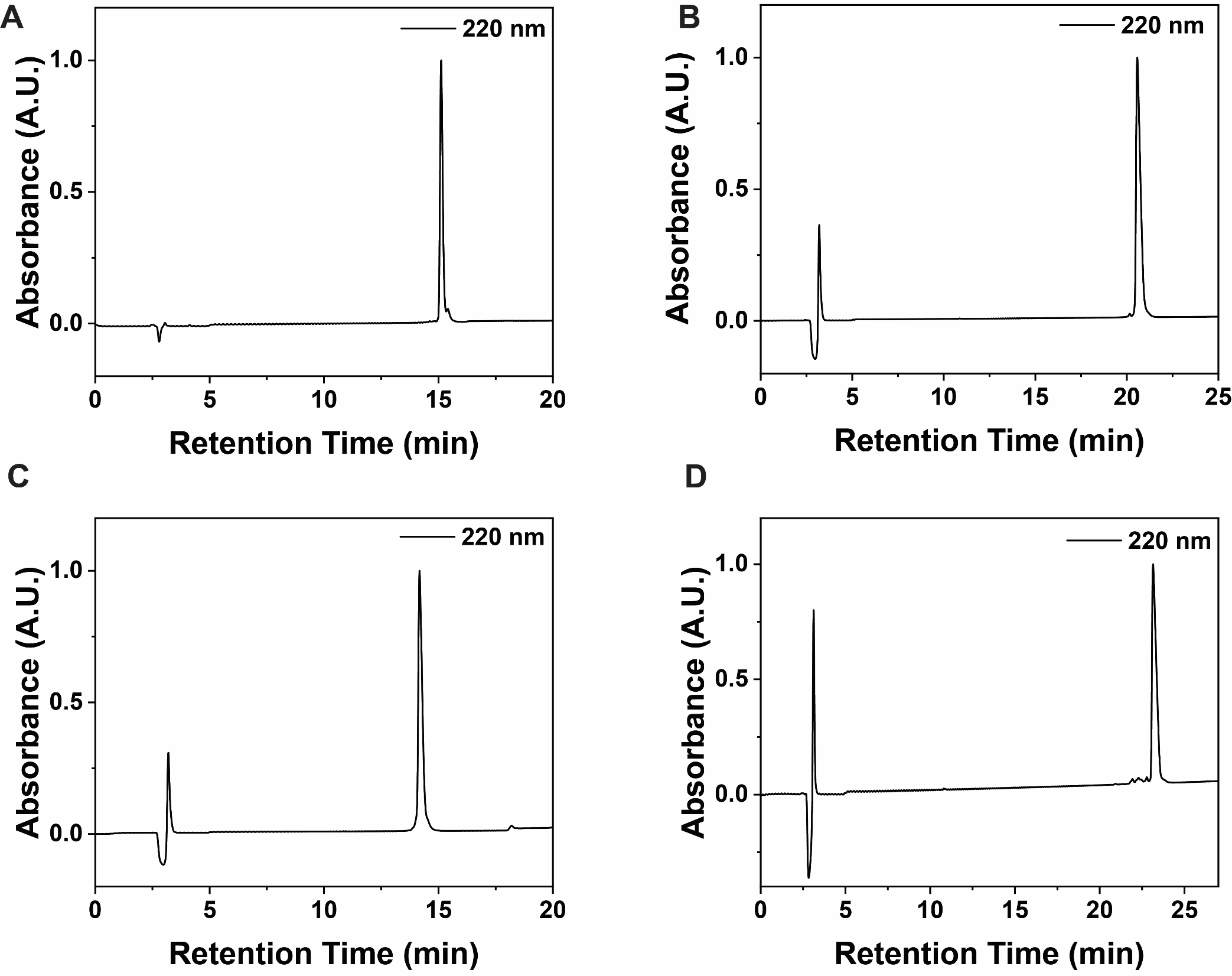

**Figure S21.** HPLC traces of purified (**A**) **B_EE_**, (**B**) **B’_KK_**, (**C**) **B_EK_**, (**D**) **B’_EK_**.

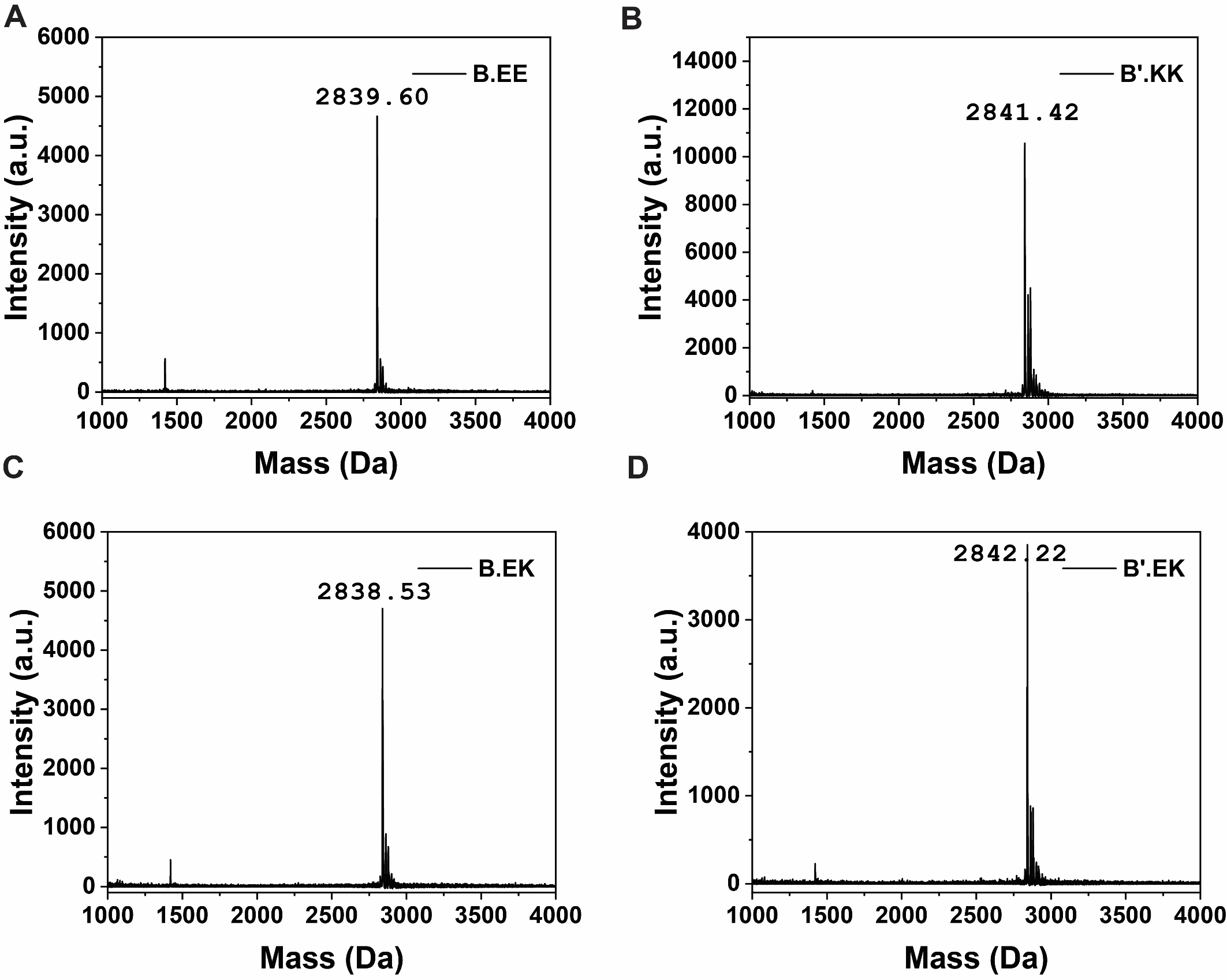

**Figure S22.** MALDI-TOF/MS spectra for purified (**A**) **B_EE_**, m/z = 2839.60 [M+H]^+^, (**B**) **B’_KK_**, m/z = 2841.42 [M+H]^+^, (**C**) **B_EK_**, m/z = 2838.53 [M+H]^+^, (**D**) **B’_EK_**, m/z = 2842.22 [M+H]^+^.

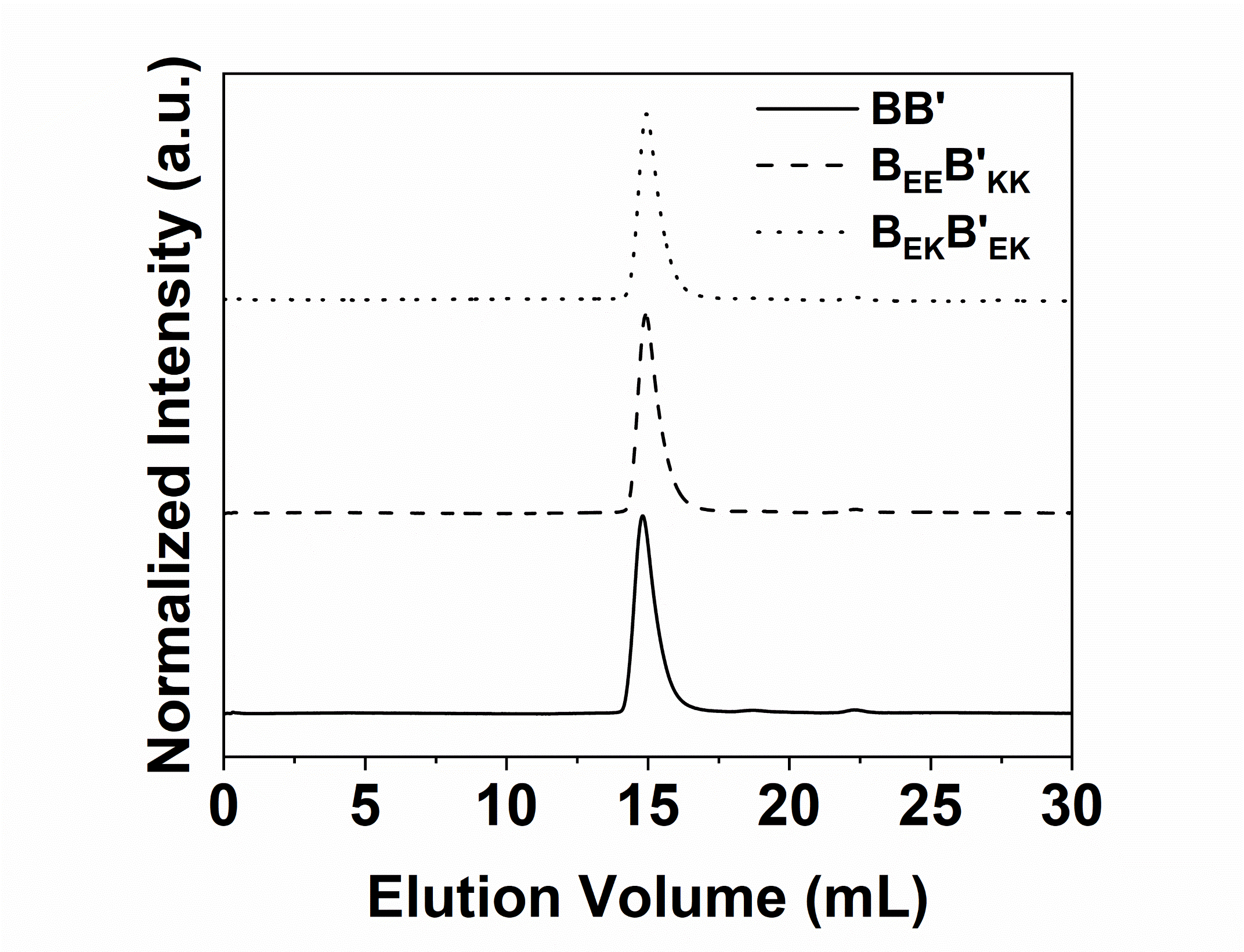

**Figure S23**. SEC profiles of **BB’**, **B_EE_B’_KK_**, and **B_EK_B’_EK_** confirming consistent oligomeric state.

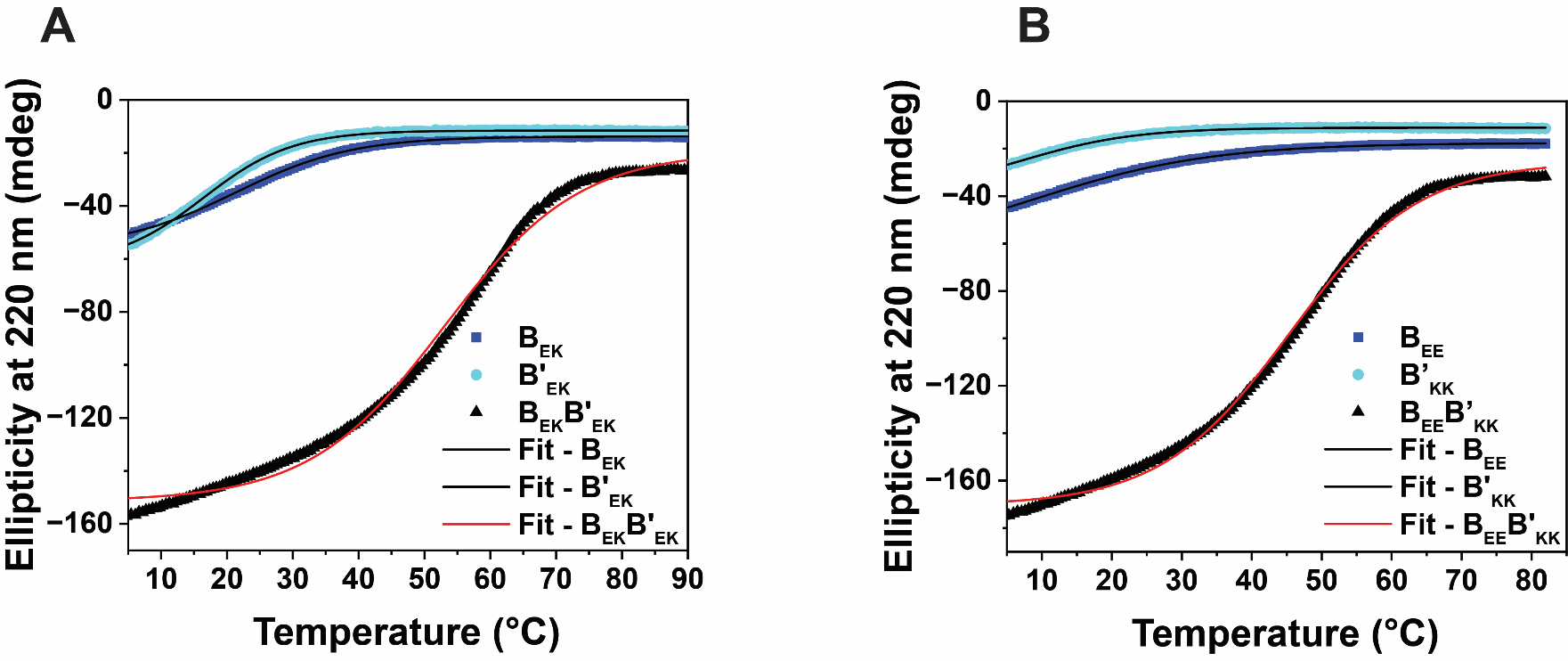

**Figure S24.** Thermal denaturation plots and their fit curves for (**A**) **B_EK_**, **B’_EK_**, and **B_EK_B’_EK_**, and (**B**) **B_EE_**, **B’_KK_**, and **B_EE_B’_KK_** and their fitted curves.

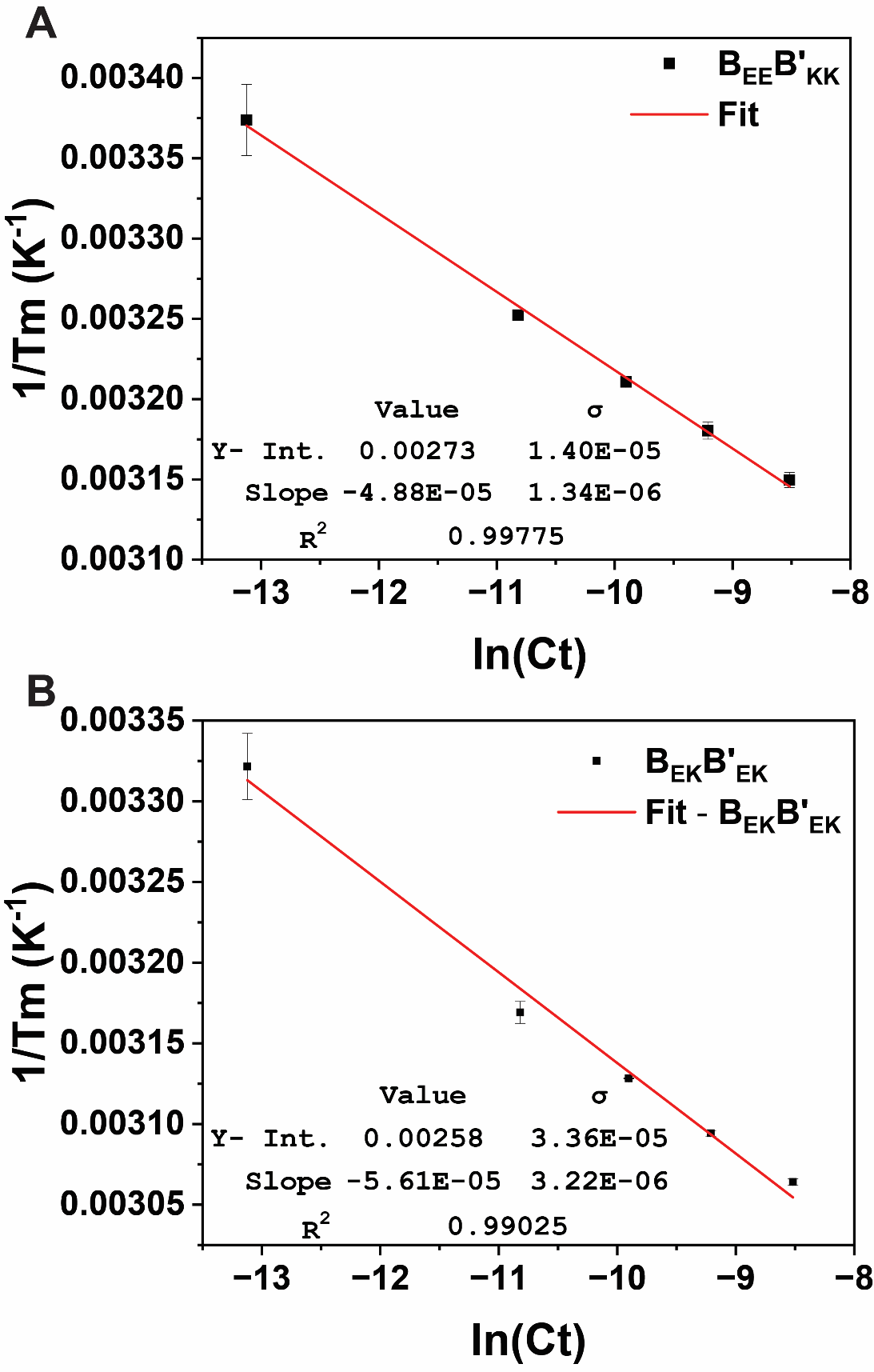

**Figure S25.** Van’t Hoff plots and analysis for (**A**) **B_EE_B’_KK_** and (**B**) **B_EK_B’_EK_**.

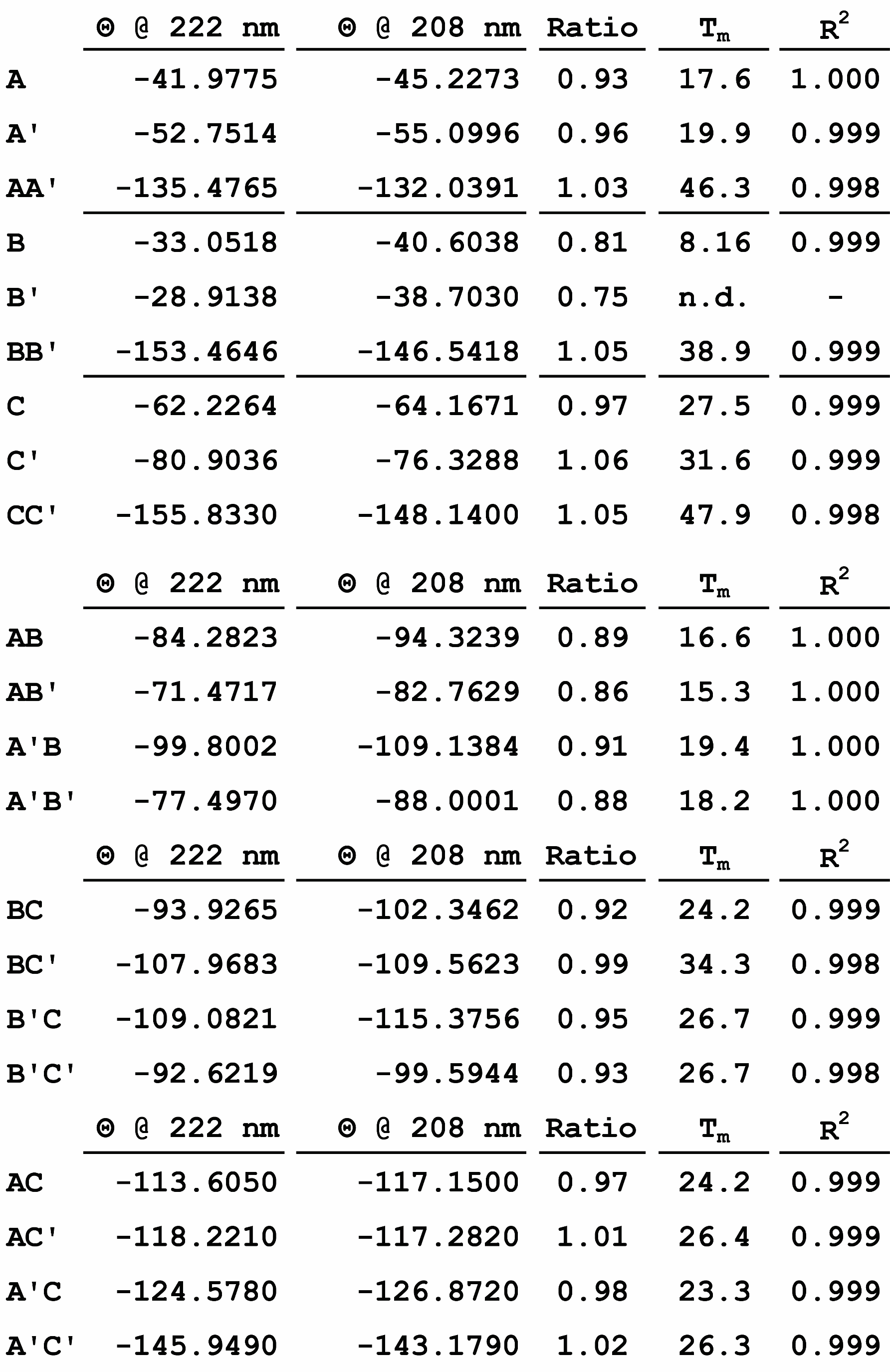

**Table S1.** Tabulated data from experimental CD data of on-target and off-target interactions (n.d. = not detected).

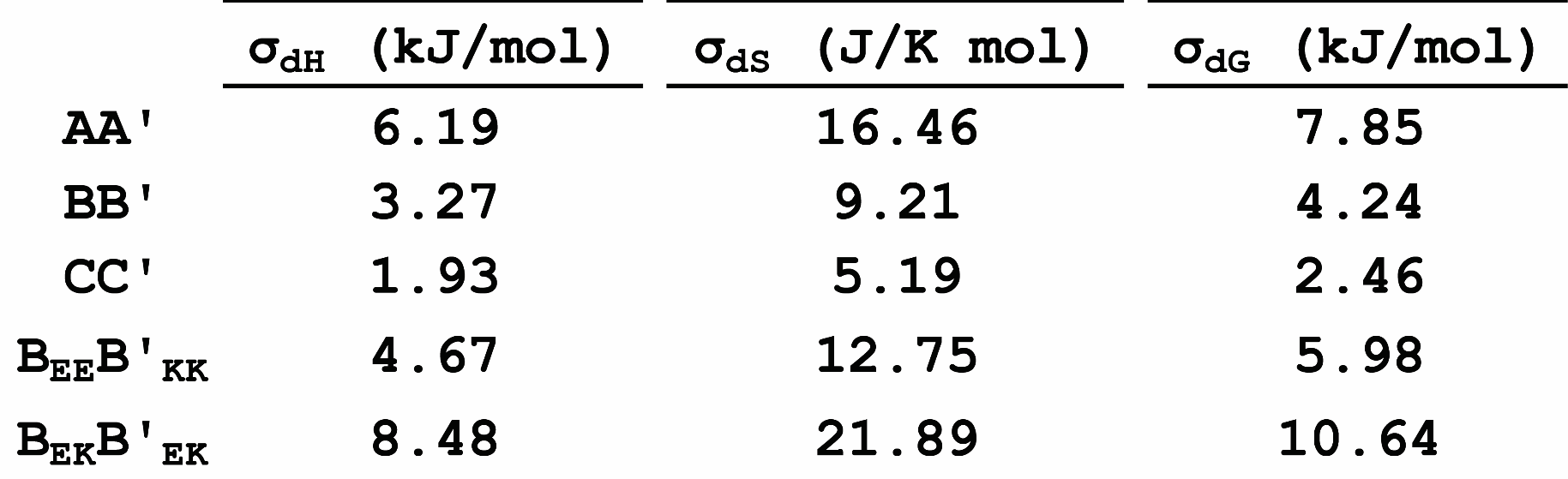

**Table S2.** Table of uncertainties calculated by propagating the error obtained from linear regression.

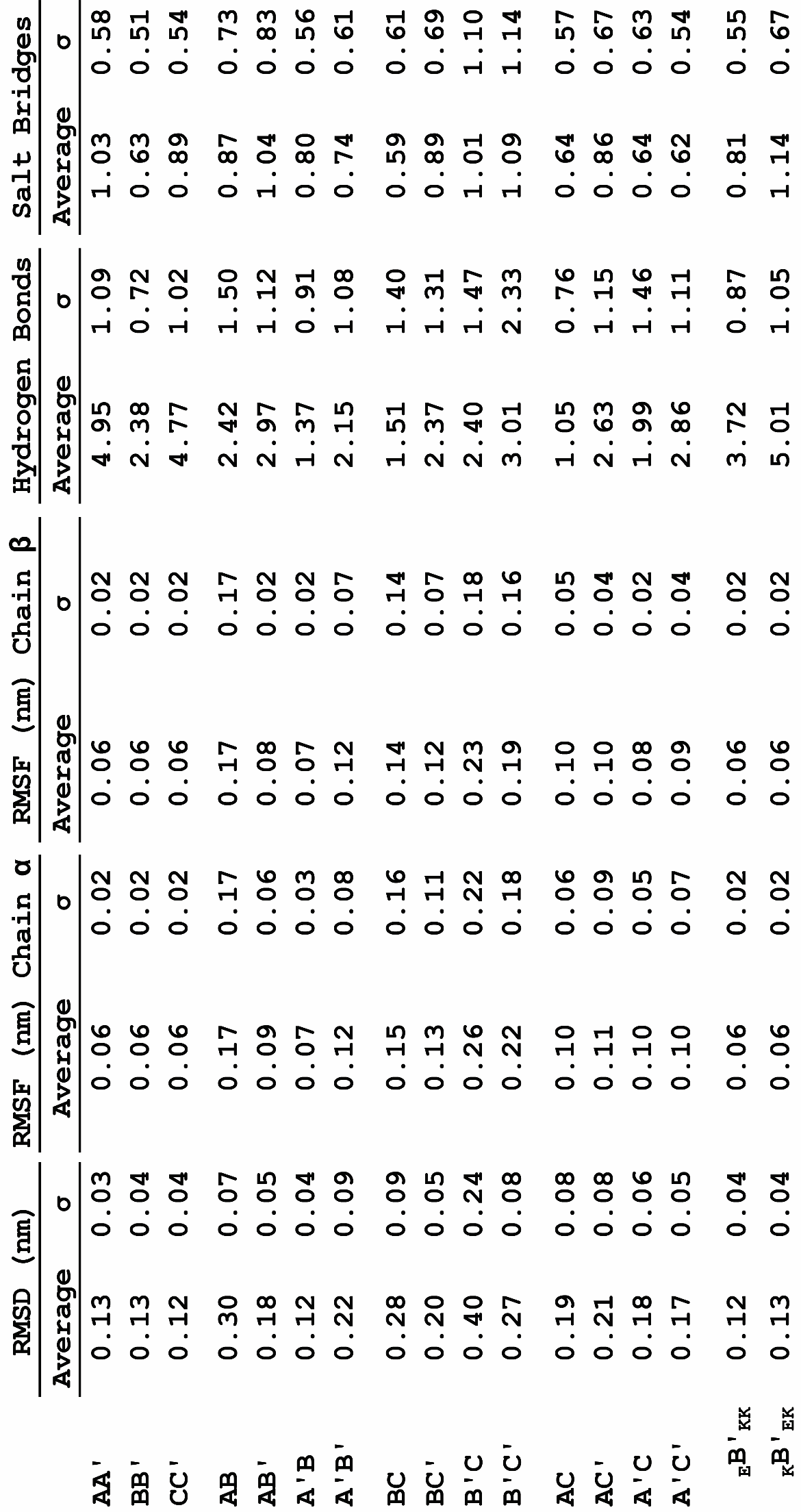

**Table S3**. Tabulated RMSD, RMSF, number of hydrogen bonds and salt-bridge contacts for the fifteen simulated CC pairs.

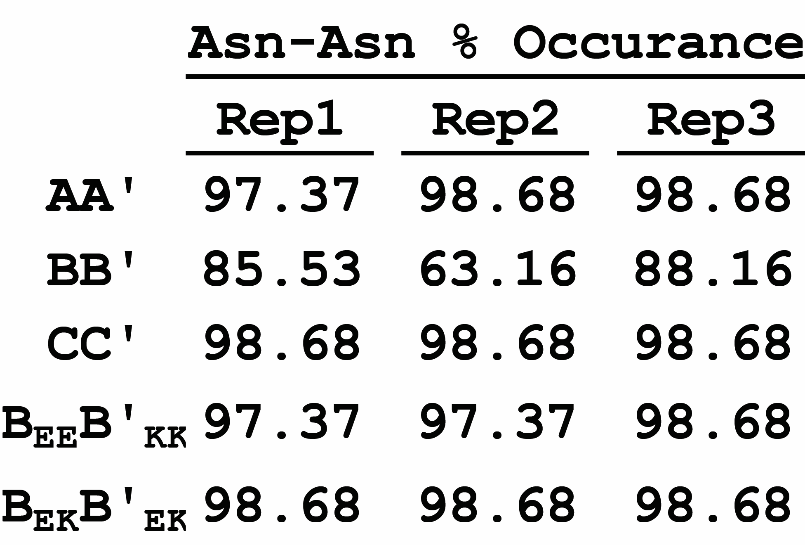

**Table S4.** Percent occurrence of Asn-Asn’ contacts for the five designed dimers during the length of the MD simulation.

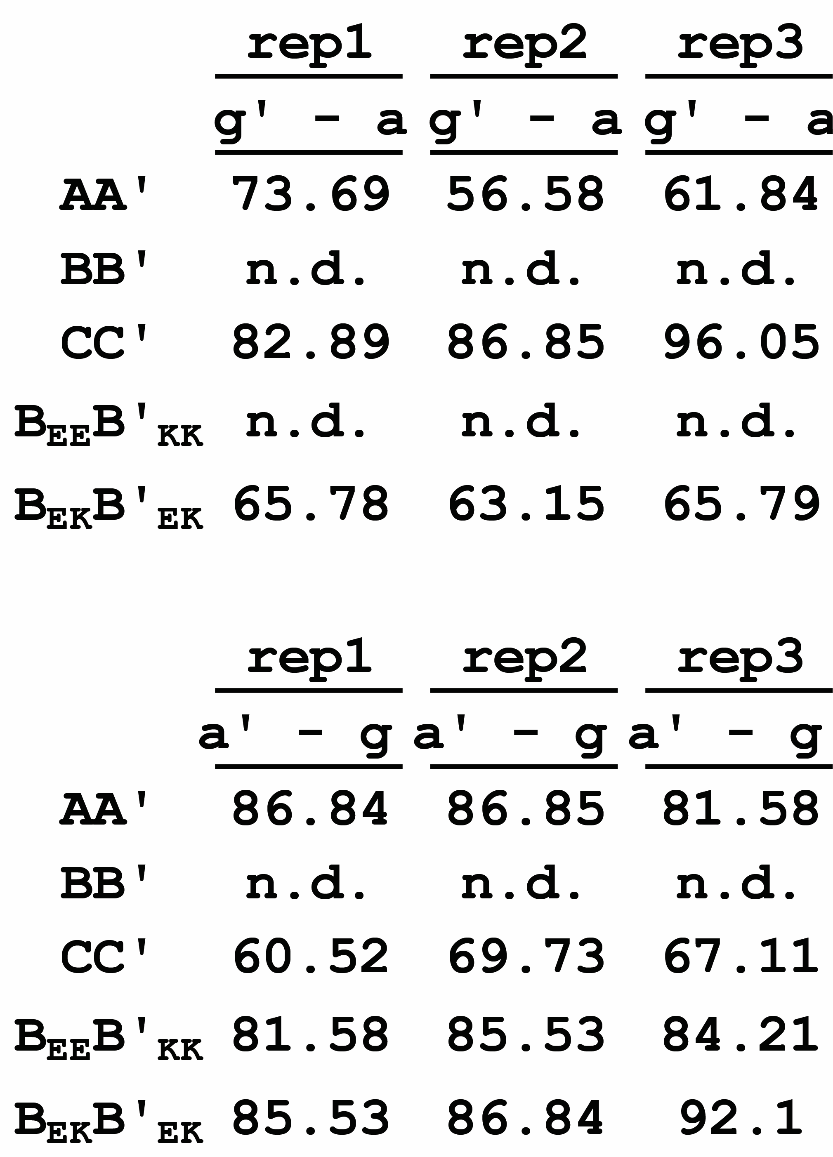

**Table S5.** Percent occurrence of Glu-Asn contacts for the five designed dimers during the length of the MD simulation.

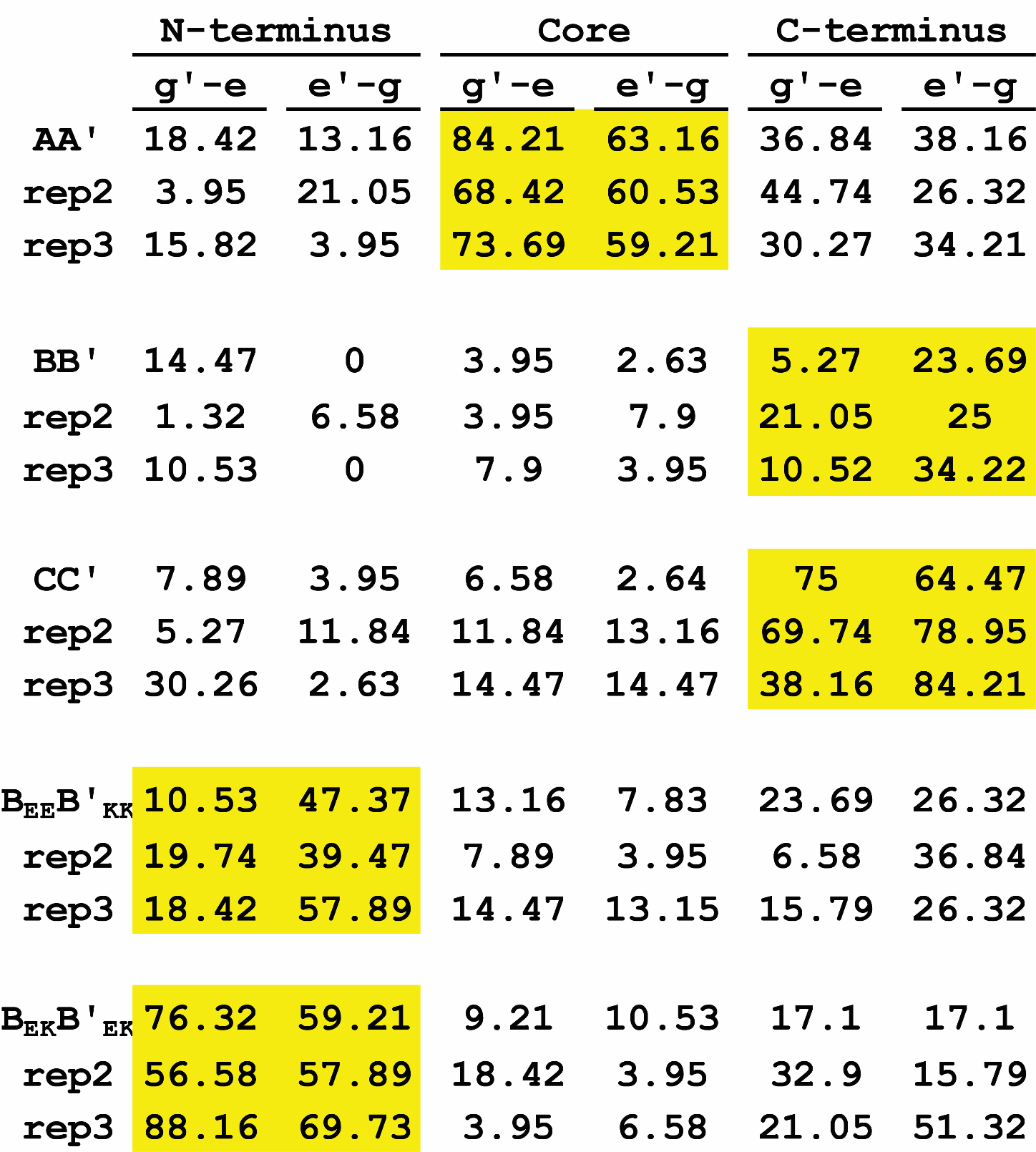

**Table S6.** Table of percent occurrence of salt-bridging interactions for the five-designed CC dimers. Highlighted portions indicate the heptad with the highest occurrence of salt bridges.

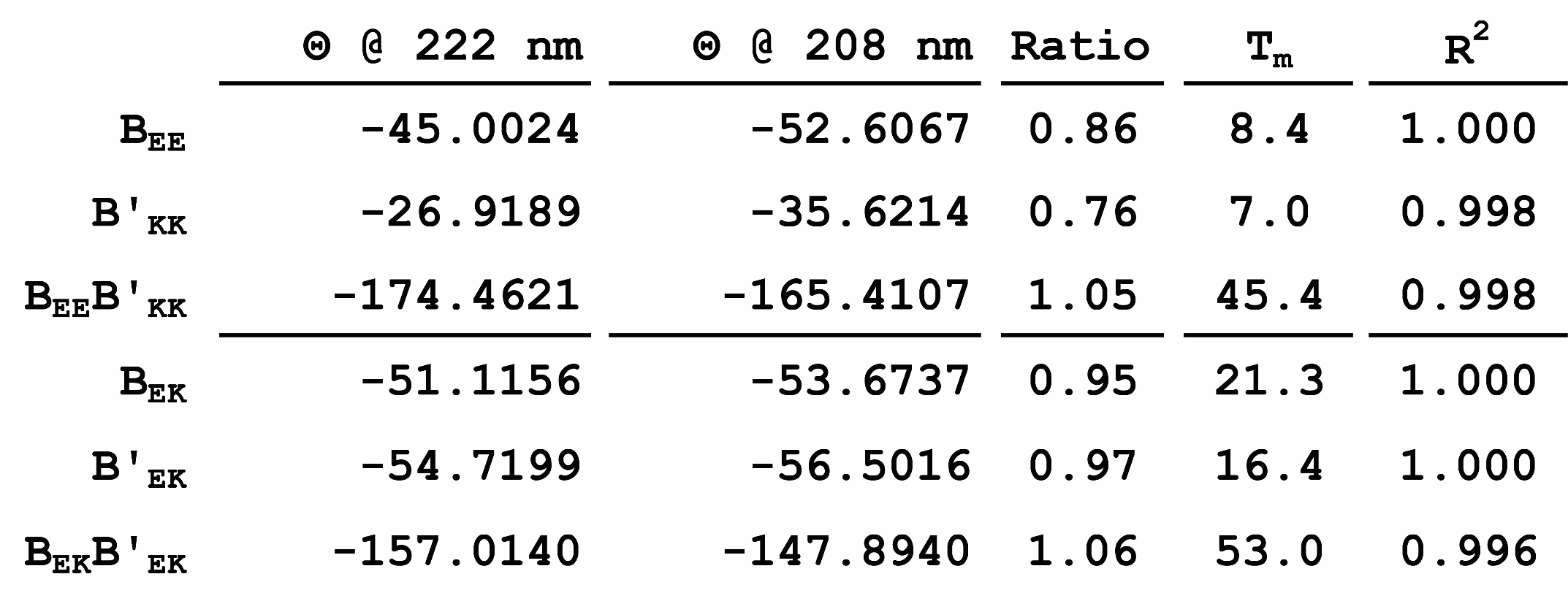

**Table S7.** Tabulated CD data for the mutated **B** and **B’** monomers and dimers. Melting temperature in degrees Celsius.
